# *In silico* experiments uncover a novel mechanism underlying mutation rate evolution in sexually reproducing populations

**DOI:** 10.1101/2021.09.25.461822

**Authors:** Andrii Rozhok, Niles Eldredge, James DeGregori

## Abstract

Natural selection is believed to universally work to lower mutation rates (MR) due to the negative impact of mutations on individual fitness. Mutator alleles can be co-selected by genetic linkage with adaptive alleles in asexual organisms. However, sexual reproduction disrupts genetic linkage, allowing selection to efficiently eradicate mutator alleles, lowering MR to the extent limited by the overall selection efficiency.

In the present paper, we apply Monte Carlo *in silico* experimentation to study MR evolution in sexually reproducing populations.

We demonstrate that both higher and lower MR can evolve depending on the mode of selection acting on adaptive phenotypic traits. We reveal a previously unreported co-selective process that determines the direction of MR evolution. We show that MR evolution is substantially influenced by multigenic inheritance of both MR and adaptive traits. Our study corroborates that MR evolution is significantly impacted by genetic drift; however, its primary source appears to be the amount of standing genetic variation, with a lesser role for population size.

Based on our study, we propose an expanded population genetics theory of MR evolution in sexually reproducing populations, with potential implications for understanding rapid adaptive speciation and related macroevolutionary patterns, as well as for human health.

**Lay summary:** Natural selection is believed to always work to lower mutation rates in sexual organisms. Here we apply a Monte Carlo model of a sexually reproducing population and demonstrate that both lower and higher mutation rates can evolve, contingent on selection acting on adaptive traits in a sexually reproducing population.

## Introduction

Mutation rates range widely among different species across the phylogenetic tree of life [1]. A number of factors have been proposed to explain the MR diversity across taxa, such as genome size [2], variation in the trade-off between the cost of efficient DNA repair and speed of replication among taxa [3,4], generation time [5,6], metabolic rate [6], and population size affecting the efficiency of selection in lowering MR (1). Many of the proposed mechanisms are based on evidence limited to certain taxonomic groups and theoretical speculation, therefore it is unclear presently how much they can be generalized across the tree of life. However, Lynch [1] does report a plausible inverse correlation between MRs and effective population sizes across a vast span of taxa, ranging from prokaryotes to mammals. Small populations appear to demonstrate higher MR proposed to result from their lower responsiveness to selection and thus an increased inefficiency of selection in lowering MR [1,7,8].

Regardless of the factors involved, the overarching currently accepted presumption, based on empirical and modeling studies, is that selection universally works to lower MR because of the negative effect of mutations on individual fitness [1,9–13]. Exceptions to this rule have been reported for asexually reproducing species in which strong genetic linkage between mutator alleles and alleles important in evolutionary adaptation in some cases leads to co-selection for mutator alleles and thus a higher MR – the “mutator hitchhiking” model [14–17]. In sexually reproducing populations, however, genetic linkage is substantially reduced by the recombination process, and such mutator hitchhiking is deemed to be the exception, not the rule. However, genetic hitchhiking might not be the only mechanism that contradicts the presumption that MRs are universally lowered by selection, and some studies do support the possibility that sexual organisms could evolve higher MRs under certain conditions [18].

The following reasoning can be proposed to argue that the sign of selection acting on MR may likely be reversible even in sexually reproducing organisms. Quantitative phenotypic traits with multigenic inheritance are often distributed in populations in terms of their expression, in a typical case following a normal distribution as shown in **Fig. 1**. Under stabilizing selection (**Fig. 1A**), the population’s mean expression of the selected trait will tend to confer the highest fitness and thus be the most frequent (although deviations from normality do occur). Lower fitness phenotypes deviate toward the tails of the phenotypic distribution and occur at lower frequencies. Based on the multigenic inheritance of MRs, the many genes shown to control DNA damage levels, and some emergent evidence [19–23], we can assume that MR is also a distributed trait in sexually reproducing populations and should vary among individuals. Evidence from recent studies already supports this view [22,24]. Hypothetically, individuals that are closer to the population phenotypic mean should come from parents that have lower MRs and inherit, among the multiple genes encoding MR, a collection of alleles that result in a relatively low MR. Correspondingly, individuals who inherit more alleles promoting higher MR should have higher chances of phenotypically deviating from the population mean across generations and ending up in one of the tails of the phenotypic distribution, as shown in **Fig. 1**. It follows then that stabilizing selection acting on the phenotype should increase the frequency of low MR alleles (by favoring mean, non-deviant phenotypes) and decrease the frequency of high MR alleles in the population. However, when selection acting on the phenotype changes to directional (**Fig. 1B**), whereby tail phenotypes are favored, higher MR alleles should undergo enrichment and low MR alleles should decrease in frequency as selection favors phenotypes that significantly deviate from the population mean.

**Fig. 1.**
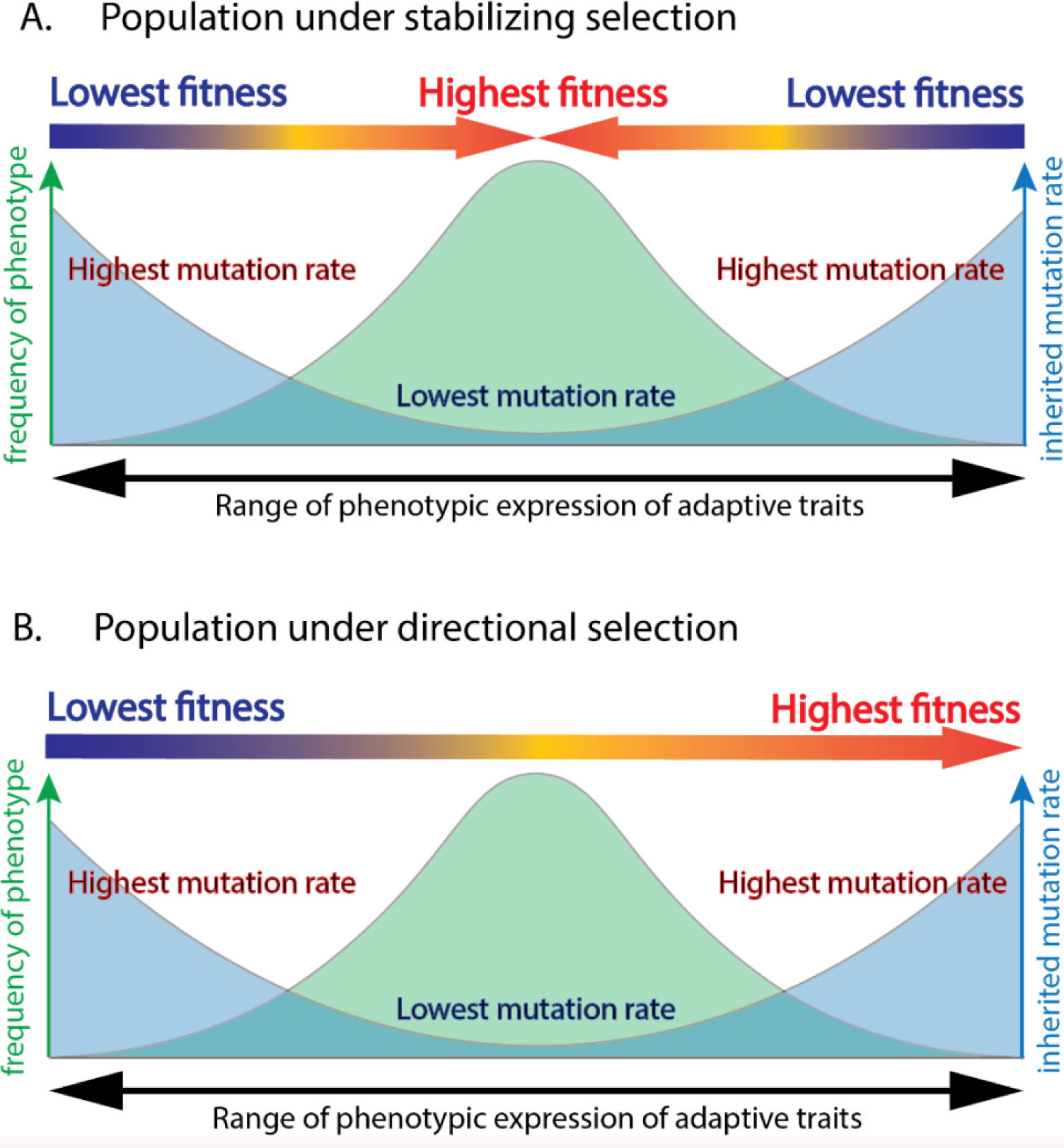
A hypothetical model for how selection regimen should impact the evolution of mutation rates. **A,** we assume a normal distribution of a quantitative distributed adaptive trait (green area) in a typical large population; as argued in the text, mutation rate (blue area) is also a quantitative distributed trait based on its highly multigenic inheritance; we assume that mutation rate is distributed so that phenotypically most deviant individuals (axis X) harbor the highest mutation rate based on the idea that such individuals are likely to inherit more alleles associated with higher mutation rate, which over many generations causes the progeny of such lineages on average to deviate phenotypically more profoundly compared to individuals whose adaptive phenotypic trait is close to the population mean; under stabilizing selection (color gradient arrows), mean phenotypes are the fittest and selected for, which should trigger co-selection for lowest mutation rates. Such co-selection for mutation rate should occur progressively across generations. **B**, when directional selection is applied to such a population, it should trigger co-selection for higher mutation rate, since the favored tailing phenotypes are more likely to harbor alleles for higher MR based on the same logic as explained in panel A.

Therefore, the overarching assumption here is that alleles of the many genes controlling MR are distributed in sexually reproducing populations such that alleles inducing lower MR are most frequent in the genomes of individuals with the least deviant phenotypes (from the optimal highest frequency / highest fitness phenotypes). The genomes of the most deviant tailing phenotypes, conversely, should be enriched in higher MR alleles that underlie the observed more frequent and more pronounced phenotypic deviation. Noteworthy, this is not to argue that MR is directly responsible on a per generation basis for creating more phenotypic variability in a population, but rather that such an effect of differential phenotypic deviation among progeny should accumulate over generations in specific sub-populational genetic lineages in an MR-dependent manner. Therefore, the existing phenotypic variation per generation should primarily come from pre-existing genetic variation and not due to mutations occurring in the current generation. The fact that MR is distributed in a population (or whenever it is distributed) corroborates that such non-homogeneities and segregation of sub-populational genetic lineages takes place and prevents homogenous distribution of traits. Based on such reasoning, therefore, it can be hypothesized that, contrary to the current conventional presumption, the direction of MR evolution in sexually reproducing populations should not always be toward lower MR, but should instead depend on the general selection regimen acting on adaptive phenotypic traits. Stabilizing selection will favor optimal non-deviant phenotypes whose genomes are enriched in lower MR alleles, and directional selection will increase the frequency of deviant tailing phenotypes enriched in higher MR alleles in the genome.

In the present study, we apply *in silico* experiments based on the Monte Carlo paradigm replicating a sexually reproducing population of organisms to test this hypothesis and look deeper into factors that are critical in the evolution of MR. Monte Carlo modeling has been extensively used in many areas of science as a tool specifically designed to recapitulate *in silico* very complex natural systems involving multiple independent processes [25]. The complexity of such systems, notably including many biological processes, at times achieves levels that are not tractable by traditional analytical approaches, and analytical solutions for such processes often do not exist. This problem is particularly apparent in stochastic processes, such as genetic change and many aspects of sexual reproduction. *In vivo* experimental approaches also are limited in their power to describe such systems due to the narrow focus of experimental manipulations, usually addressing only particular aspects of a complex system at a time. Monte Carlo modeling, therefore, is part of a very limited set of tools that can provide insights into how complex systems operate as a whole and provide a guidance for further *in vivo* experimental studies.

Our results reveal a previously unreported mechanism modulating the sign of selection acting on MR that is consistent with our hypothesis shown in **Fig. 1**. Our results demonstrate theoretical evidence that selection for higher MR is possible and may be common in sexually reproducing organisms. These results have broad implications, from understanding evolutionary change under rapidly changing environments to human health.

## Materials and Methods

### Software

The model code (**Supplement 1** “Model Matlab code”; further referred to as “Code”) was created and *in silico* experimentation performed using the Matlab software (The MathWorks Inc., Natick, Massachusetts). Variables from the Code are italicized in the explanation below for convenience. **Fig. 2** illustrates major model steps (refer to the Code for explanation of each step).

**Fig. 2.**
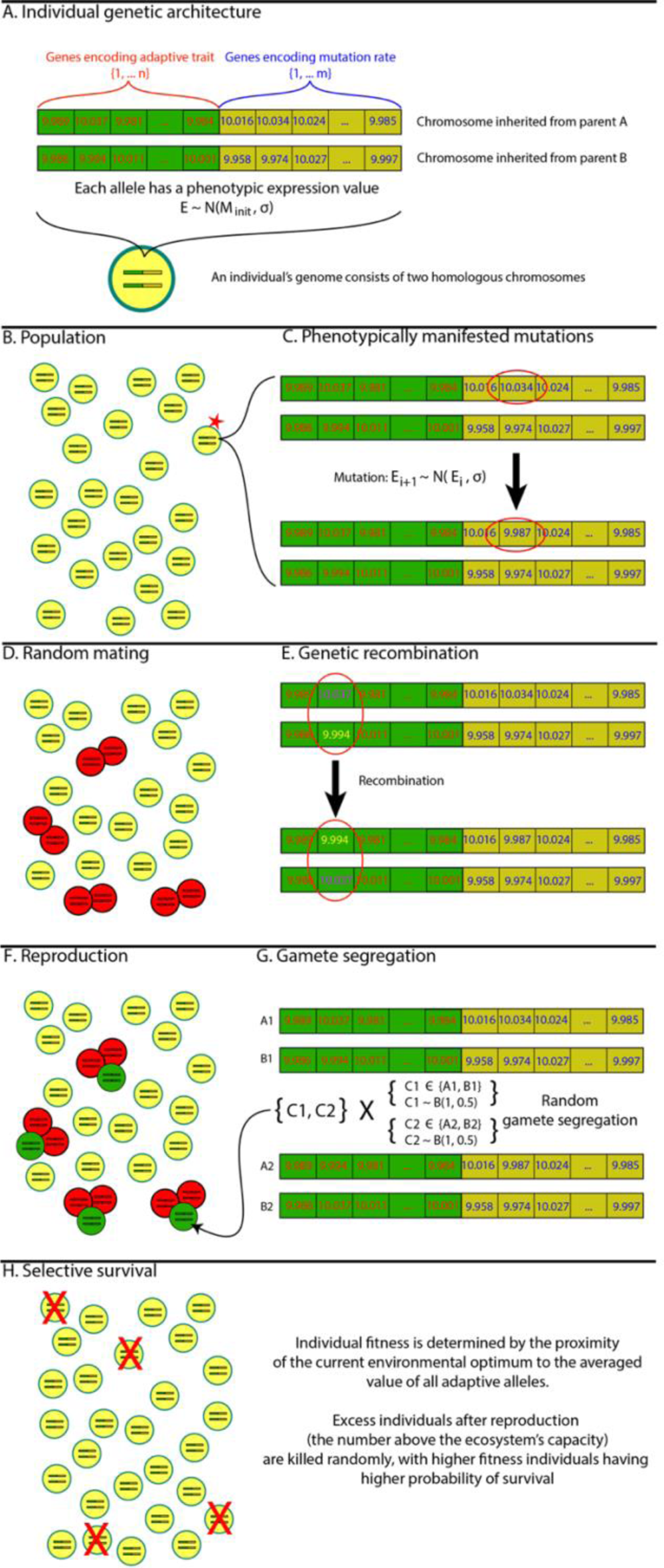
Major model steps. **A**, An individual’s genome consists of two homologous chromosomes each inherited from one of the individual’s parents. Each chromosome consists of two sets of genes: a) genes that encode adaptive phenotypic traits (left side in green) and b) those that encode MR (right side in yellow). Each gene has an expression level (E) that in the initial population is generated from a normal distribution with the mean M (initial population optimum of 10) and a standard deviation representing variance of inheritance *σ* (= variance of functional mutation effects). **B**, The population consists of sexually reproducing organisms and has a maximum population size imposed by the ecosystem’s capacity. **C**, Functional mutations occur at the current individual’s MR determined by the averaged expression of all alleles encoding MR multiplied by a fixed transformation coefficient common to all individuals. Functional mutations alter the expression value of the affected gene such that the gene’s altered expression level is drawn from a normal distribution with the pre-mutation expression serving as the mean and a standard deviation *σ*. **D**, A subset of individuals (based on the reproduction frequency preset in the model) randomly mate for reproduction. **E**, Genetic recombination occurs at a preset rate per gene per simulation step and is modeled as homologous alleles swapping their respective chromosomes. **F**, Each reproducing pair of individuals (red circles in D and F) produce a single progeny (green circles) per mating, with each individual reproducing multiple times throughout the simulation. **G**, Gamete segregation is modeled by using Bernoulli trials such that each chromosome in each parent has a 50% chance of being transmitted to the progeny. **H**, After reproduction, progeny are added to the population and selective elimination of individuals follows by applying Bernoulli trials for each individual such that an individual’s chance of survival depends on the individual’s fitness. An individual’s fitness is determined by averaging the expression of all adaptive alleles and comparing it to the current environmental optimum (proximity means higher fitness). The overall frequencies of selective mortality are such that the remaining population approaches the ecosystem’s capacity.

### The adopted paradigm of mutation rate (MR), distribution of phenotypic effects (DPhE) and distribution of fitness effects (DFE) of mutations for modeling evolution

The model simulates a generic case of a population of organisms, each containing a diploid genome, each haplotype of which consists of one chromosome carrying genes encoding MR and genes encoding the simulated adaptive trait. We ignore the rest of the genome as unrelated to the relationship studied. Cells reproduce by mating and producing a single progeny per reproduction which inherits one random chromosome from each parent.

Genetic and epigenetic changes (mutations) are diverse and have multifarious effects. Part of mutations are silent, and other have physiological consequences for cells and organism. In this study, we ignore silent mutations to replicate a basic generic evolutionary process whereby selection only operates on mutations that have immediate individual phenotypic effects. The entire set of phenotype-altering mutations is integrated into a single normal distribution, such that after each simulation iteration each simulated gene undergoes a binomial trial with probability *P* for whether a phenotype altering mutation has occurred in it and, if so, the mutation’s phenotypic effect is drawn based on the parameters on the underlying normal distribution. Therefore, the mutation rate adopted in the model is not any actual mutation rate or its approximation, but instead is a generic rate of phenotype-altering mutations. Because the rate of phenotype-altering mutations for any given genomic architecture (species) is linearly related to its overall mutation rate, it is safe to assume that changes in the rate of phenotype-altering mutations will be informative on changes in the actual mutation rate. Noteworthy that even the rates of phenotype-altering mutations that we model are not an approximation of real rates from any species, but rather a generic value whose evolutionary response (change) in response to adaptive evolutionary processes is the focus of the present study. For simplicity, we therefore refer to this rate as mutation rate (MR). The reason we adopt the normal distribution for mutation phenotypic effects is, again, to reflect a simple generic case of a typical distribution of continuously distributed phenotypic traits, known as the Shelford’s law [26]and widely represented in natural populations.

Functional mutations do not have a direct impact on organismal fitness and thus selection. Functional mutations alter phenotype. Fitness, as manifested in allele frequency alterations over generations, arises when selection is applied to a given phenotype. Therefore, as a random process mutations produce a distribution of phenotypic effects (DPhE), and when selection is applied to a population it generates a distribution of mutation fitness effect (DFE) as a function of DPhE and selection. Therefore, unlike DPhE which is mostly determined by the genome architecture of a species, DFE is a very dynamic distribution that depends on what selection regimen is applied to a given diversity of phenotypes. We therefore only explicitly model selection regimens and the distribution of phenotypes (as the main determinants of the shape of DFE in natural populations), while DFE itself naturally evolves over a simulation run without control from the modeler and can only be measured retrospectively after the run. Such architecture recapitulates typical relationships between mutations, phenotype, selection, and fitness in natural evolution.

### Model algorithm

#### Pre-simulation steps

**1:** model parameters are initialized (Code lines 16-29):

*maxTime* – number of simulation iterations whereby model state update is done after each iteration. Default 10,000.

*popsize* – maximal population size cap as the maximum number of individuals allowed to exist in the simulated population, recapitulating the concept of ecosystem’s carrying capacity (ECC); if the population becomes larger than this cap, the overcrowding becomes an additional factor of mortality weighed by individual relative fitness, which recapitulates intra-specific competition limited by ECC. Default 10,000.

*nMutGenes* – the number of genes in the simulated genome affecting mutation rate. Default 10.

*nAdaptGenes* – the number of genes in the simulated genome encoding the simulated distributed adaptive trait. Default 10.

*initAllele* – initial phenotypic manifestation of the simulated genes, being a number representing the initial phene/phenotype produced by a gene. This number, being generic and unrelated to any specific value in natural genomes, is applied for simplicity to both – genes encoding MR and those encoding the simulated adaptive trait. Therefore, each gene for MR and for adaptive trait starts from the default phenotypic value V ∼ N(*initAllele*, δ), the mean being 10 by default, which represents simply an initial phenotypic expression of a gene contributing to a quantitative multigenic distributed trait (which both MR and the simulated adaptive trait are). The resulting trait’s (both MR and adaptive trait) phenotype is then calculated as the average phenotypic expression of all the genes that encode the trait. We assume a simple generic case that all genes contribute to the encoded trait equally, and since the phenotype of 10 or any such number is generic, there is no difference in how exactly the encoding genes interact (averaging, additive or multiplicative effects) to produce the net phenotype, since the goal of the study is to track changes, not absolute values, in MR as a response to selection acting on a adaptive trait.

*initMutRate* – a service variable used elsewhere in the code to ensure that the standard deviation of mutation DPhE is proportional to MR. This proportionality is necessary to model only functional mutations as a normally distributed DPhE without directly modeling MR by assuming that at higher MR the total frequency of functional mutations will also be higher and thus their cumulative effect on a continuously distributed phenotypic trait will likewise be larger. Default 0.1.

*inhVar* – initial standard deviation of DPhE determining the magnitude of the phenotypic effects of functional mutations. Default 0.05.

*percentReproducing* – percent of the simulated population involved in reproduction at each simulation iteration. Default 0.05 (5%).

*recombinRate* – a change that a given gene recombines (switches chromosomes) per simulation step. Default 0.05 (5%).

*envChange* – a service variable determining the rate of change of the highest fitness phenotype (goal phenotype) under directional selection. Directional selection results from environmental change and works in a way that over time the best phenotypic expression (highest fitness) of an adaptive continuously distributed trait shifts into the selected direction. Default 0.001.

*costOfMutRate* – the somatic cost associated with mutations which is manifested in risks, such as inherited phenotypic aberrations, cancer etc., modelled as a low impact individual fitness penalty for having a higher MR.

**2**: setting mode of selection (Code lines 34-35, **Supplement 2**). Selection is modelled in such a way that proximity of an individual’s expression of the adaptive trait to the best fitness (phenotypic goal) value increases the individual’s fitness relative to others. For example, the initial goal phenotype in all simulations is 10. Individuals whose overall expression of the adaptive trait is closer to 10 have higher chances of survival per each simulation step. Under stabilizing selection regimen (Code line 34), the best fitness phenotype (goal phenotype) always remains 10 (set by modeler), so mutant deviants have reduced fitness. Under directional selection (Code line 35), the goal phenotype is set to be moving over the course of the simulation in the selected direction in *envChange* increments (simulation of a major natural environmental change that triggers directional selection), favoring individuals whose phenotype progressively deviates over generations into the selected direction. Fitness (preferential survival) is realized as a result of ECC capped intra-specific competition described above (see explanation for the *popsize* variable).

**3**: creating initial population (Code lines 40-45, 50-55). An initial population of *popsize* individuals is created, each individual is assigned a diploid genome with each haplotype being represented with a chromosome consisting of *nMutGenes* and *nAdaptGenes* of genes encoding MR and the adaptive trait, respectively. Each gene of the genome in each individual (both for MR and adaptive trait) is set to its initial phenotypic value V ∼ N(*initAllele*, δ). As a result of this setup, an initial population *of popsize* individuals is established that demonstrates a normally distributed variation of individual MR and adaptive phenotype, following the Shelford’s law assumption [26].

#### Simulation run

The model run (Code lines 69-215) is a Monte Carlo stochastic iteration over the simulated time of 1 through *maxTime* (10,000) simulation steps with the following events at each step:

1. MUTATION (Code lines 74-91): each simulated gene in each individual undergoes a binomial trial for whether a phenotypically manifesting mutation has occurred in the gene. As explained above, we do not simulate mutations or MR directly, but instead the model applies a certain low probability that such a mutation has occurred and, if true, assigns a slight phenotypic change to the encoded phenotype drawn randomly from the DPhE distribution of a given individual. Such mutations in genes encoding the adaptive trait lead to changes in the phenotypic value of the adaptive trait, while in genes encoding MR lead to changes in the probability of such mutations in genes of a given individual.
2. RECOMBINATION (Code lines 99-124): the simulated genetic recombination (crossing over) is modelled as a small probability for a given gene to switch chromosomes. Natural crossing over typically occurs as an exchange of entire chromosome arms, thus often many genes switch chromosomes together (known as genetic linkage), with genes located closer to each other having higher chances to recombine together. This process creates linkage disequilibrium, whereby shuffling of alleles among haplotypes in the population deviates from idealized theoretical randomness. However, in order to produce a generic idealized population for the purpose of a theoretical study, we do not model genetic linkage, and therefore each allele in the simulated population is recombined independently of others and is thus randomly shuffled among haplotypes. Therefore, for the purpose of critical assessment of the results of the present study, it is important note that genetic linkage was absent in the modelled population.
3. REPRODUCTION (Code lines 133-171): the population consists of sexually nondifferentiated individuals who can randomly mate with any other individual. At each simulation step, every individual undergoes a binomial trial with probability 0.05 for whether a given individual will reproduce at this step (see explanation above for *percentReproducing*). Approximately 5% of population is randomly chosen in this way and randomly paired for mating. A mating pair produces a single progeny (in order to equalize reproductive success) which inherits one of each parent’s chromosomes with 50% probability (simulated random gamete segregation). As with recombination, we chose such a sexless architecture in order to achieve the effect of a generic ideal population whereby the effective “sex ratio” is always close to 50/50 %, since any individual can potentially mate with any other. Factors, such as genetic linkage mentioned above, sex ratio, variation in reproductive success, age structure, migration are known to significantly affect the effective population size, Ne, relative to census population size (55), making *N_e_* typically much smaller than census population size. We eliminated these phenomena from our ideal panmictic (all individuals are potential partners) population with an effective sex ratio close to 50/50 %, no age structure, equalized reproductive success per mating, and absent migration and genetic linkage, making thus the *N_e_* maximally close to population census size. Such an architecture makes the population census size a close proxy to *Ne*, which can be used directly to compare the effects of *N_e_* on MR evolution with effects of selection. Given the comparative nature of our study within one and the same model architecture used in all simulations, population census size may seem to be a good relative proxy for *N_e_* even in a more realistic (non-ideal) population genetic setup. However, the ratio of *N_e_* relative to population census size is subject to evolution in non-ideal populations (27). Therefore, an ideal panmictic population model is better suited for a theoretical study such as the one presented here.
4. SURVIVAL (Code lines 179-195): as described above, individual fitness in the model was primarily determined by the proximity of the phenotypic expression of the simulated adaptive trait to the best fitness value imposed by the applied selection regimen. Under stabilizing selection individuals with the value of the adaptive trait closest to the initial goal of 10, which remained stable throughout the simulation run, were the fittest. Under directional selection the best phenotype goal was moving in the direction of the selected tail of the phenotypic distribution (environmental shift), therefore the individuals that were the fittest were constantly replaced over generations with new phenotypes that evolved their simulated adaptive trait to keep up with the constantly changing phenotypic goal (simulated process of adaptation). Individual fitness was also slightly negatively impacted by higher MR, reflecting the cost of somatic mutations. Each individual at each step was tried in a binomial trial for whether it survives the present simulation step, with the probability that weighed in relative individual fitness and the surplus of the population after each reproduction cycle over the ECC-imposed cap size so that on average the fittest survived best and the population after each survival trial was brought back (approximately) to its ECC cap (10,000 individuals by default). Such an architecture left only one factor impacting mortality – relative individual fitness, recapitulating thus the intra-specific competition-related mortality in natural populations. Other mortality causes, such as predation, diseases and aging were excluded from the model architecture for the same purpose as genetic and reproductive irregularities – in order to replicate an ideal panmictic population in which all processes related to the evolution of a sexually reproducing population were appropriately randomized and thus excluded from the study as potential confounding variables.

## Results

### Selection acting on adaptive traits impacts the evolution of mutation rate

We first proceeded with testing our overarching hypothesis that the mode of selection acting on an adaptive trait will impact the evolution of MR. For Fig.2, modeling employs our default parameter values. See **Model algorithm. Pre-simulation steps** in **Materials and Methods** for a full description of parameters and their justification, as well as model design outlined in **Fig. 2**. As shown in **Fig. 3A-B**, under stabilizing selection the expression of the adaptive trait remains relatively constant and close to the initial population mean, while under directional selection it responds to selection toward increased expression. This measurement served as a control in order to ensure that the selection regimens indeed worked as expected in their effect on the adaptive trait. Another control, ensuring that the population numbers stayed at the imposed ecosystem’s capacity is shown in **Fig. 3C-D**, and confirms that the imposed population size maximum worked as expected. **Fig. 3E-F** demonstrates that the initial distribution of MR did not differ between the stabilizing and directional selection setups (t-test; p=0.49; respective means equaled 10.004 and 9.993). **Fig. 3G-H** shows that at the end of the simulation the distribution of MRs significantly differs between selection regimens and provides support for our primary hypothesis that directional selection acting on adaptive traits alters the mode of evolution of MR (Wilcoxon rank sum test; p<0.001; means: 6.81 and 16.35). Wilcoxon ranks sum test will be used in testing all further results regarding the final MR. As we anticipated based on our model shown in **Fig. 1**, under stabilizing selection MR evolves toward lower values (mean=6.81), while directional selection leads to higher MRs (mean=16.35) in our experiment. Notably, directional selection creates initial transient drops in population size by, perhaps, imposing higher initial rates in mortality when the environment is changing and altering the fitness peak. During later stages of evolution, the population appears to catch up with the change and restores is maximum size. This pattern will be observed throughout the remaining experiments.

**Fig. 3.**
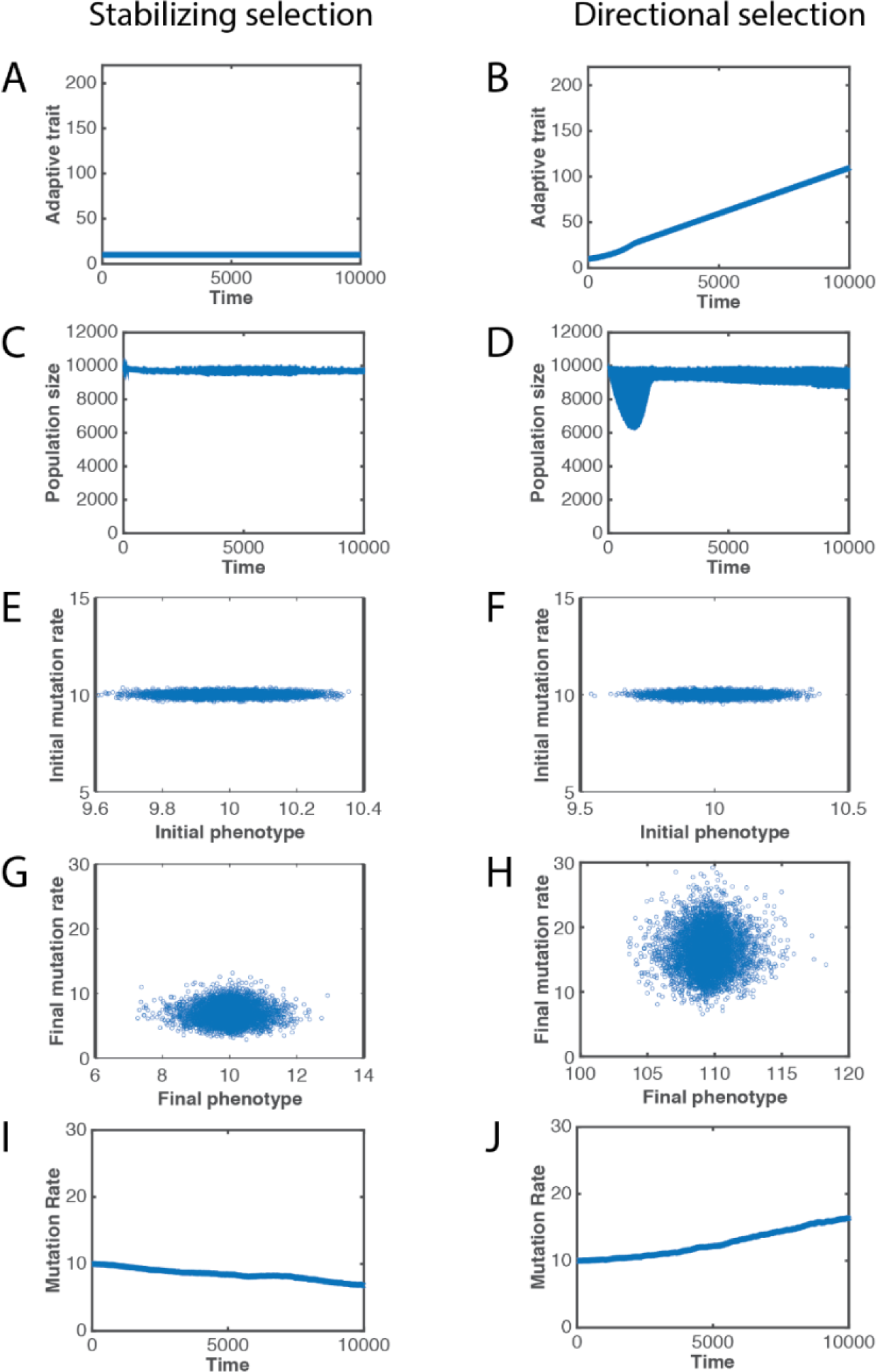
The effect of selection acting on an adaptive trait on the evolution of mutation rate. Left panels represent evolutionary processes when the adaptive trait is under stabilizing selection. Right panels represent evolutionary processes when the adaptive trait is under directional selection toward increased expression. **A-B,** Average adaptive trait expression (in arbitrary units) throughout the simulation time (in simulation updates). **C-D,** The dynamics of population (in the number of individuals) throughout the simulation time. **E-F** The distribution of MR (in arbitrary units) relative to the distribution of initial expression of the adaptive trait; each circle represents one individual. **G-H,** The distribution of MR relative to the distribution of adaptive trait expression at the end of the simulation. **I-J,** The evolution of the average MR in the population throughout the simulation time.

### Stronger directional selection increases the observed differences in MR evolution

We further aimed to explore whether the regimen of selection acting on the adaptive trait was at least part of the causes that lead to the differences in MR evolution observed in **Fig. 3G-H,I-G**. We therefore increased the strength of directional selection by 30% relative to that applied in the experiment shown in **Fig. 3**. Henceforth, we will only show the distributions of MRs at the end of each simulation, while the dynamics of the control processes will be shown in supplementary figures. Increasing the strength of directional selection leads to a statistically significant increase in MR (**Fig. 4A-B**; p<0.0001; means: 16.35 and 20.85), providing an indication that directional selection acting on the adaptive trait is at least partially responsible for the observed differences in the evolution of MR. As expected, the adaptive trait (the X-axes in **Fig. 4A-B**) also demonstrated a dramatic increase in expression in response to stronger directional selection. Population dynamics and other controls are shown in **Supplement 4a** (note a deeper initial drop in population size in response to a faster environmental change).

**Fig. 4.**
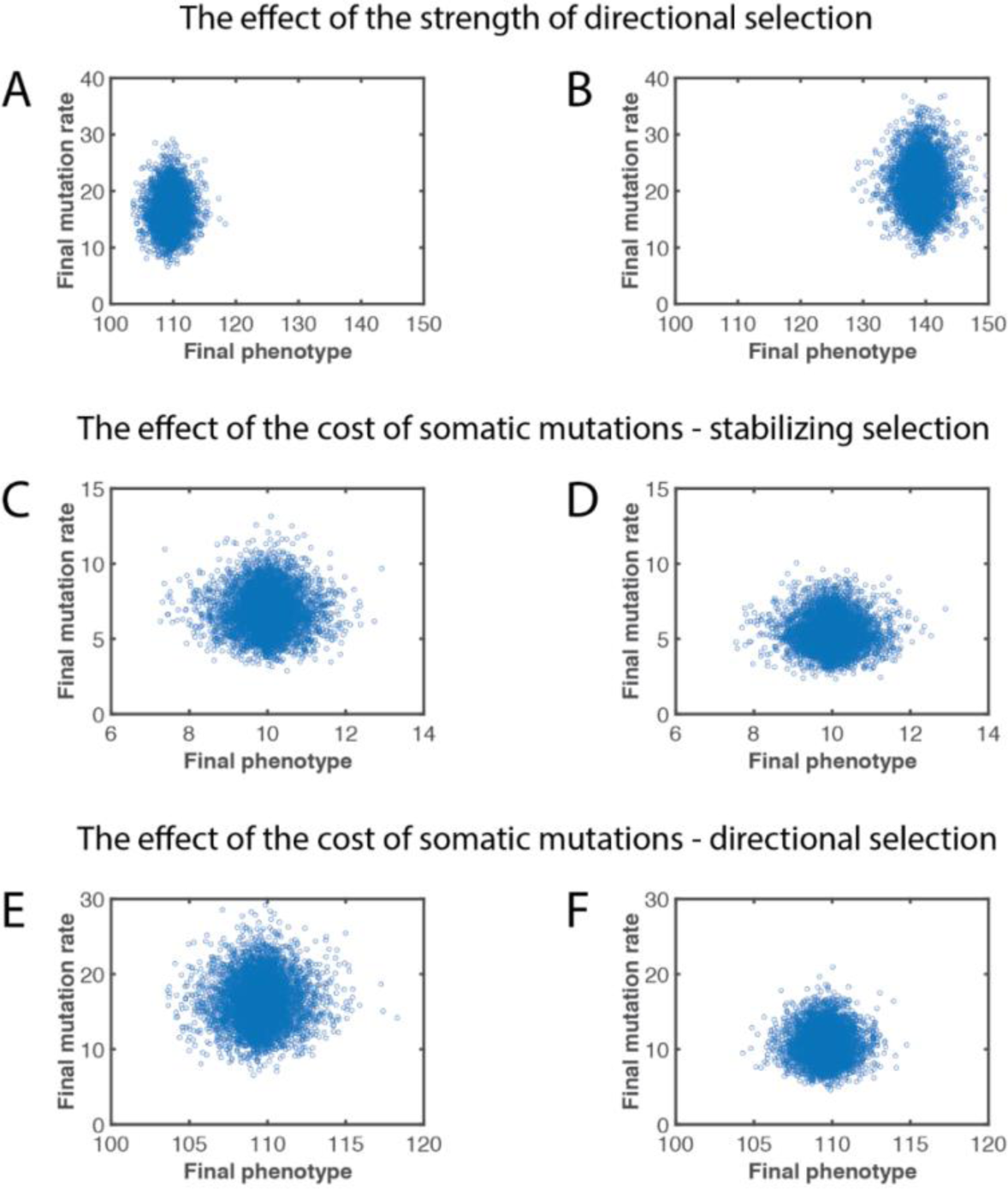
The effect of the strength of directional selection and the cost of somatic mutations on MR evolution. **A,** MR evolution under the default (parameter value 0.001) strength of directional selection acting on the adaptive trait. **B,** MR evolution under increased (parameter value 0.0013) strength of directional selection acting on the adaptive trait. **C,** MR evolution with the default cost of somatic mutations (parameter value 0.001) and stabilizing selection acting on the adaptive trait. **D,** MR evolution with an elevated cost of somatic mutations (parameter value 0.1) and stabilizing selection acting on the adaptive trait. **E,** MR evolution with the default cost of somatic mutations (parameter value 0.001) and directional selection acting on the adaptive trait. **F,** MR evolution with an elevated cost of somatic mutations (parameter value 0.1) and directional selection acting on the adaptive trait.

### The cost of somatic mutations alters MR evolution under both stabilizing and directional selection

Theoretically, the evolution of MR shown above should be affected by the cost of somatic mutations, with the higher cost promoting selection for lower MR. In order to elucidate the role of the cost of somatic mutations, we increased the cost by 100-fold (by increasing 100-fold the individual fitness penalty for having a higher MR) under both stabilizing and directional selection acting on the adaptive trait. Under stabilizing selection, a higher cost of somatic mutations promoted stronger selection for lower MR, resulting in lower MR values (**Fig. 4C-D**; p<0.001; means: 6.81 and 5.4). **Fig. 4E-F** demonstrates that a higher cost of somatic mutations has a similar effect under directional selection by buffering the evolution of higher MR and leading to lower MR values at the end of simulation (p<0.001; means: 16.35 and 10.6). Even with the lower MR, the final selected phenotype was similar for the two conditions, despite insignificant evolution of MR with the higher cost of somatic mutations. Control processes are shown in **Supplements 4b** and **3c**, correspondingly. Therefore, the combined results shown in **Fig. 4C-F** are consistent with the idea that the cost of somatic mutations is a modulator of MR evolution, and a higher cost promotes selection for lower MR regardless of the regimen of selection acting on adaptive traits.

### Multigenic inheritance of the adaptive trait has a significant impact on the evolution of mutation rate

If the hypothetical mechanism proposed in **Fig. 1** is correct, we should also expect that selection acting on a monogenically inherited adaptive trait will result in a different pattern of MR evolution compared to a multigenic adaptive trait. In a multigenic quantitative distributed trait encoded by many genes roughly equally contributing to the net phenotype, on average more mutations should be required to achieve the same phenotypic change, as each gene only influences the phenotype partially. Therefore, under directional selection acting on a multigenic adaptive trait, we should expect a tendency toward a higher resulting MR, since selection will favor individuals that demonstrate a higher ability to deviate into the selected phenotypic tail. The ability of stabilizing selection acting on the adaptive trait to lower MR should, likewise, be buffered by multigenic inheritance of the adaptive trait, since a monogenic trait requires lower MR to be changed in its phenotypic manifestation whereas selection acts to maintain proximity of the trait to the selected optimum. In addition, under both selection regimens, the encoding of the adaptive trait across multiple genes will hamper segregation of alleles by recombination, the latter being the main cause of independent selection acting on MR and adaptive traits and thus the believed universal action of selection to lower MR. Data shown in **Fig. 5** agree with such expectations. As can be seen in **Fig. 5A**, under stabilizing selection acting on a multigenic (10 genes) adaptive trait, evolution toward lower MR is observed (See also **Fig. 3**). If the adaptive trait is monogenic (**Fig. 5B**), a significantly lower MR evolves (p<0.001; means: 8.81 and 1.42), consistent with the idea that a monogenic trait requires fewer mutations to be altered, therefore selection can lower MR further. As expected for a monogenic trait, a much narrower distribution of phenotypes is observed (**Fig. 5B**). Multigenic traits require more mutations per the same amount of phenotypic change, and therefore higher MR is tolerated whereby the phenotypic expression of the adaptive trait remains stable. Controls are shown in **Supplement 5a**. Theoretically, if each mutation has less effect on the phenotype, we should see a reverse trend toward the higher MR under monogenic inheritance of the adaptive trait, since in such a scenario more mutations will have less effect on the trait. We made the inherited DPhE of mutations narrower (the parameter value was lowered from the standard 0.05 to 0.01 under stabilizing selection and 0.018 under directional; the latter value is higher, since 0.01 under all other parameters unchanged rendered the population non-responsive to directional selection by providing too little phenotypic variance and the population died out early in the simulation unable to adapt to the changing environment). **Fig. 5C** demonstrates that a narrower DPhE indeed restores higher values of the evolved MR (mean=8.23; all three experiments differ between each other with p<0.0001), consistent with the idea of a higher tolerance of a population to higher MR when mutations have smaller effects on the phenotype. Notice that, as expected, panels with lower MR also demonstrate lower phenotypic variance (X-axis; **Fig. 5A-C** in standard deviations: σ=0.567, σ=0.095, σ=0.18). **Fig. 5D-F** demonstrates a very similar pattern under directional selection acting on the adaptive trait, with monogenic inheritance of the adaptive trait leading to lower MR. Narrowing the DPhE leads to a higher final MR that is still lower than the initial MR. All the three experiments showed significant differences (p<0.001) of the means and variances (controls for Fig. **5D-E** are in **Supplement 5b**, for **Fig. 5C-F** in **Supplements 5c** and **5d**, respectively). Indeed, with monogenic inheritance, we fail to observe any evolution towards increased MR under the standard DPhE (**Fig. 5E**). As before, note that the final phenotype is similar despite differences in MR. The results shown in **Fig. 5**, therefore, corroborate the idea that the number of genes encoding adaptive traits that are under various regimens of selection significantly impacts MR evolution.

**Fig. 5.**
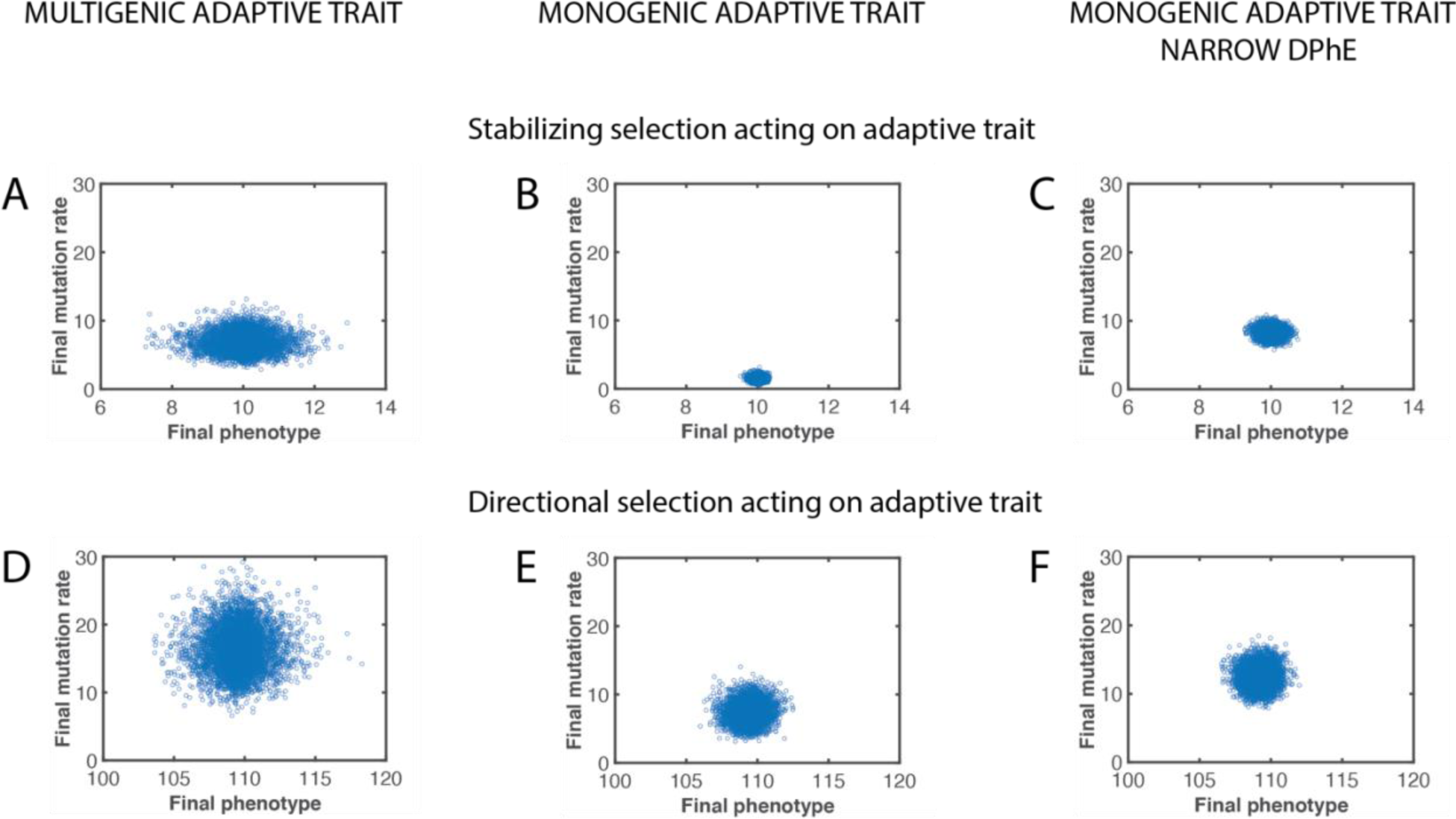
The effect of multigenic inheritance of the adaptive trait on MR evolution. Left column: multigenic adaptive trait, middle column: monogenic adaptive trait, right column: monogenic adaptive trait with a narrow distribution of phenotypic effects (DPhE) of mutations. **A-C,** The effect of multigenic inheritance of the adaptive trait under *stabilizing* selection acting on the adaptive trait. **D-F,** The effect of multigenic inheritance of the adaptive trait under *directional* selection acting on the adaptive trait.

### Multigenic inheritance of mutation rate buffers the effect of selection

We further hypothesized that the number of genes encoding MR should also impact MR evolution. If theoretically MR was encoded by one gene, the gene would be better visible to selection given its direct association with the phenotypic expression of MR. Since selection acts on genes indirectly via phenotypic traits, multigenic inheritance should make each particular gene less visible to selection because each gene only partially contributes to the trait. As shown in **Fig. 6A-B** (controls in **Supplement 6a**), under stabilizing selection acting on the adaptive trait, selection is more efficient in lowering a monogenic MR compared to multigenic (p<0.001; means: 6.81 for multigenic MR and 1.26 for monogenic MR), revealing thus the hypothesized buffering effect of multigenic inheritance of MR on the ability of selection to lower MR under stabilizing selection. **Fig. 6C-D** (controls in **Supplement 6b**) also reveals a similar effect when the adaptive trait is under directional selection (p<0.001; means: 16.35 and 4.12) – indeed, even with directional selection, there is potent selection for *lower* MR when monogenetically encoded. The latter pattern can be explained by a direct visibility of the single gene underlying the MR trait to selection whereby the cost of MR triggers selection for a lower MR. In addition, for monogenically encoded MR an allele that increases MR can easily be segregated away from the adaptive trait by recombination, which is hampered for multigenic MR. Similar to adaptive traits, multigenic encoding of MR ensures lower visibility of each gene to selection, and alleles for higher MR accumulate in the population following the principle shown in **Fig. 1**.

**Fig. 6.**
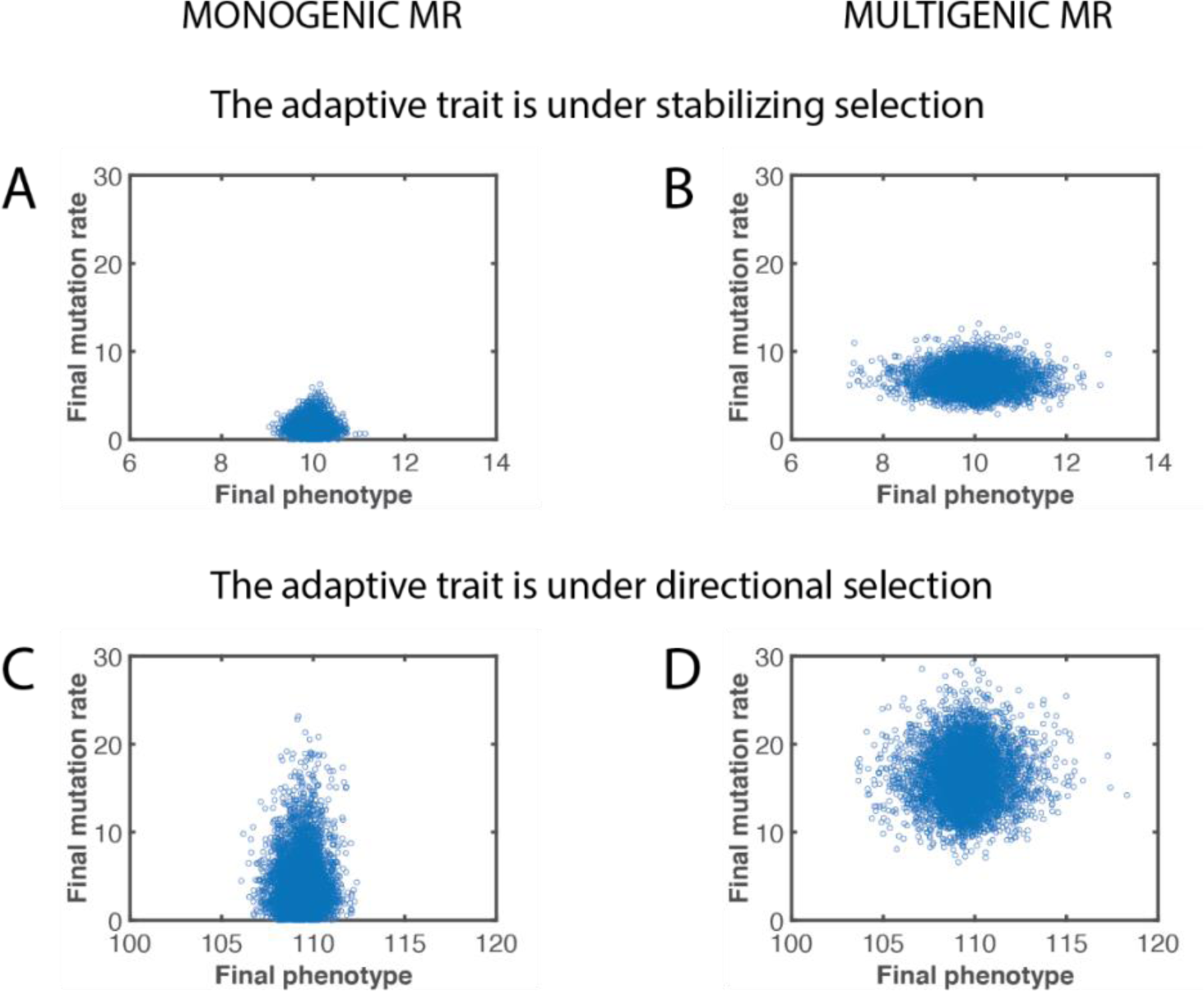
The effect of multigenic inheritance of MR impacts its evolution. **A,** Evolution of monogenically (1 gene) encoded MR under *stabilizing* selection acting on the adaptive trait. **B,** Evolution of multigenically (10 genes) encoded MR under *stabilizing* selection acting on the adaptive trait. **C,** Evolution of monogenically (1 gene) encoded MR under *directional* selection acting on the adaptive trait. **D,** Evolution of multigenically (10 genes) encoded MR under *directional* selection acting on the adaptive trait.

### Genetic recombination buffers MR evolution under both stabilizing and directional selection acting on the adaptive trait

Clonally reproducing organisms, such as bacteria, have been shown to evolve higher MR when mutator alleles are genetically linked with traits providing higher fitness [14,16]. In sexually reproducing organisms, genetic recombination is capable of effectively separating mutator alleles from adaptive alleles, which is the primary reason it is believed that selection for higher MR is not possible in populations with sexual reproduction [13,27,28]. Our results presented so far indicate that selection for higher MR is still possible even in the presence of genetic recombination. However, genetic recombination should be a factor which at least buffers selection acting on MR. We compared the evolution of MR in our simulated population in the standard setting versus a population with absent recombination, all other processes left intact (e.g., MR and the adaptive trait are each encoded by 10 genes). As can be seen in **Fig. 7A-B** (controls in **Supplement 7a**), genetic recombination buffers selection for lower MR under stabilizing selection acting on the adaptive trait (p<0.001; means: 6.81 with and 2.78 without recombination). Comparison of **Fig. 7A** and **Fig. 7B** also reveals a significant contribution of genetic recombination to the phenotypic diversity evident from the much larger variance of the phenotypic expression of the adaptive trait in the presence of genetic recombination, as expected. A similar buffering effect is visible under directional selection (p<0.001; means: 16.35 and 21.0), whereby recombination (**Fig. 7C**) obstructs the accumulation of high MR alleles in the population relative to the setting with no recombination (**Fig. 7D**). Controls for **Fig. 7C-D** in **Supplement 7b**. Therefore, our results demonstrate consistency with the general population genetics theory and indicate that genetic recombination is an important factor in MR evolution.

**Fig. 7.**
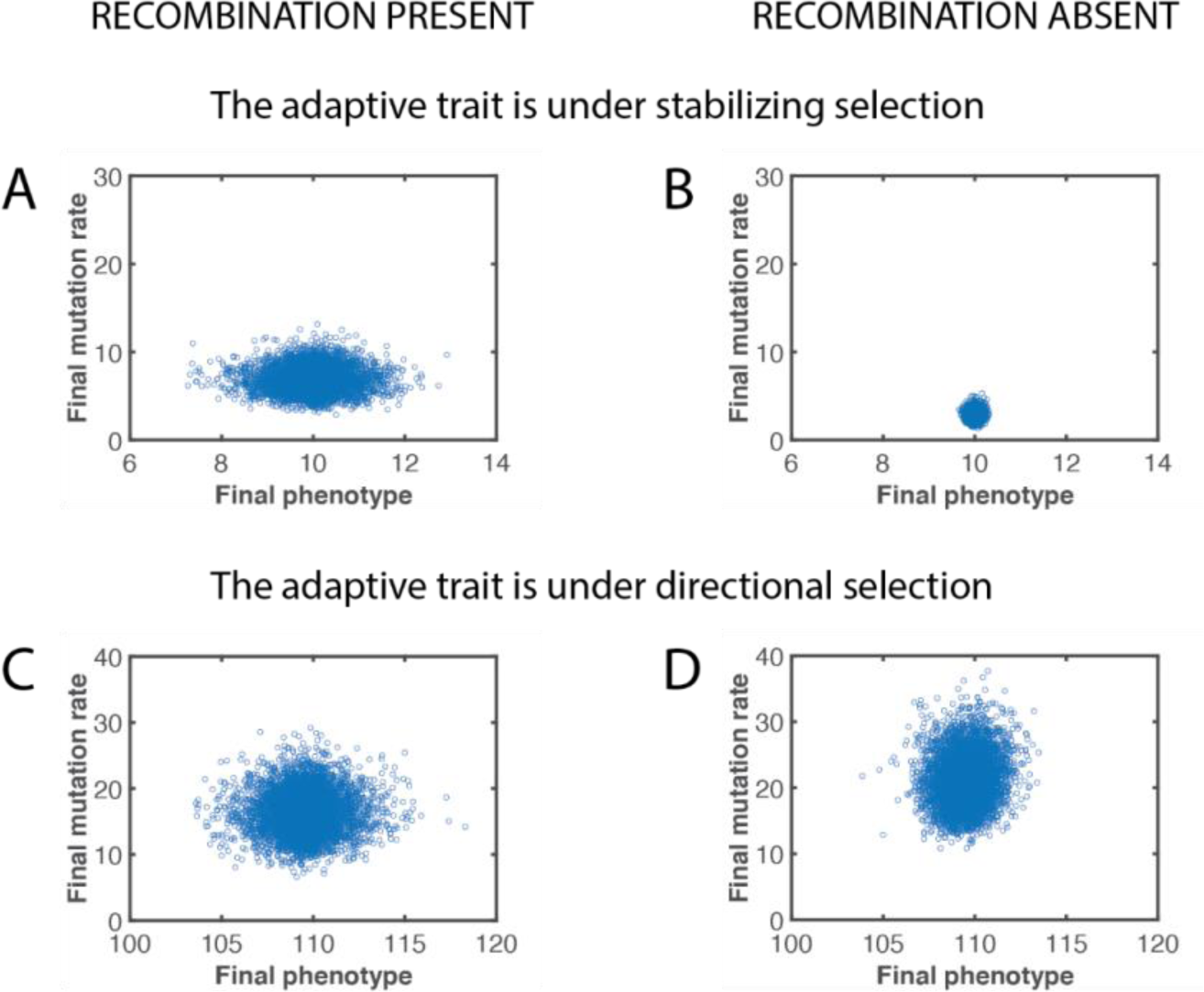
The effect of genetic recombination on MR evolution. **A,** MR evolution in presence of genetic recombination under *stabilizing* selection acting on the adaptive trait. **B,** MR evolution in the absence of genetic recombination under *stabilizing* selection acting on the adaptive trait. **C,** MR evolution in presence of genetic recombination under *directional* selection acting on the adaptive trait. **D,** MR evolution in the absence of genetic recombination under *directional* selection acting on the adaptive trait.

### Genetic drift significantly impacts the evolution of mutation rates

Lynch [1] proposed the idea that if selection works to lower MR, then MR distribution might be caused by different efficiency of selection in different species due to differences in the power of genetic drift. He demonstrated a correlation between MR and effective population size (*N_e_*) across taxa. As theoretically substantiated by Wright and Fisher [29,30], the power of genetic drift is inversely related with *N_e_*, and thus low *N_e_*’s amplify genetic drift and thus reduce the efficiency of selection. However, presumptions underlying the Wright-Fisher model do not include selection, mutation or migration, and therefore the Wright-Fisher model quantifies the pure effect of *N_e_* only. In real populations that are under selection and the effects of other population processes, *N_e_* is not the only source of genetic drift. Based on the theory of selection coefficients [31], the balance between the efficiency of selection and the amount of genetic drift is also influenced by the amount of genetic variation in a population. Simply explained, selection is not effective if all phenotypes in a population are the same. The efficiency of selection will increase, and the power of drift will decrease with increasing phenotypic (and thus the underlying genotypic) differences among individual phenotypes in a population. Populations with more standing variation will thus respond to selection more efficiently [32–34].

We therefore further tested the effect of genetic drift on MR evolution by manipulating the amount of drift based on both these parameters - population size and genetic variance. There are a number of approaches to calculate the *N_e_* of real populations. Our modelled population provides a close approximation of a panmictic ideal population and therefore its *N_e_* approximates its census size *N_c_*, which we will use for testing the effects of *N_e_* ranging 500 to 5,000,000. In order to test the effects of genetic variation, we manipulated the population parameter *inhVar* (see code in **Supplement 1**) which represents the standard deviation of the distribution of phenotypic effects (DPhE) of mutations. Both, our adaptive trait and MR are normally distributed quantitative traits, therefore variance of DPhE directly affects the amount of phenotypic variation at the beginning before selection (standing variation) and variation generated in the process of selection.

As shown in **Fig. 8**, both tested factors affecting the power of genetic drift show effects on MR evolution. However, the effect of population size (in our case approximating *N_e_*) is surprisingly modest across a range of four orders of magnitude and is most evident in populations between 500 and 50,000 individuals. On the other hand, narrow DPhEs (smaller standard deviations of inherited phenotypic variation) greatly increase genetic drift, leading to higher MR under stabilizing selection. Basically, if mutations have smaller phenotypic effects, a proportionally higher MR is needed to achieve the same amount of deviation from the phenotypic optima for traits. At least in the experiments in our model, genetic variation appears to be the dominant factor controlling genetic drift.

**Fig. 8.**
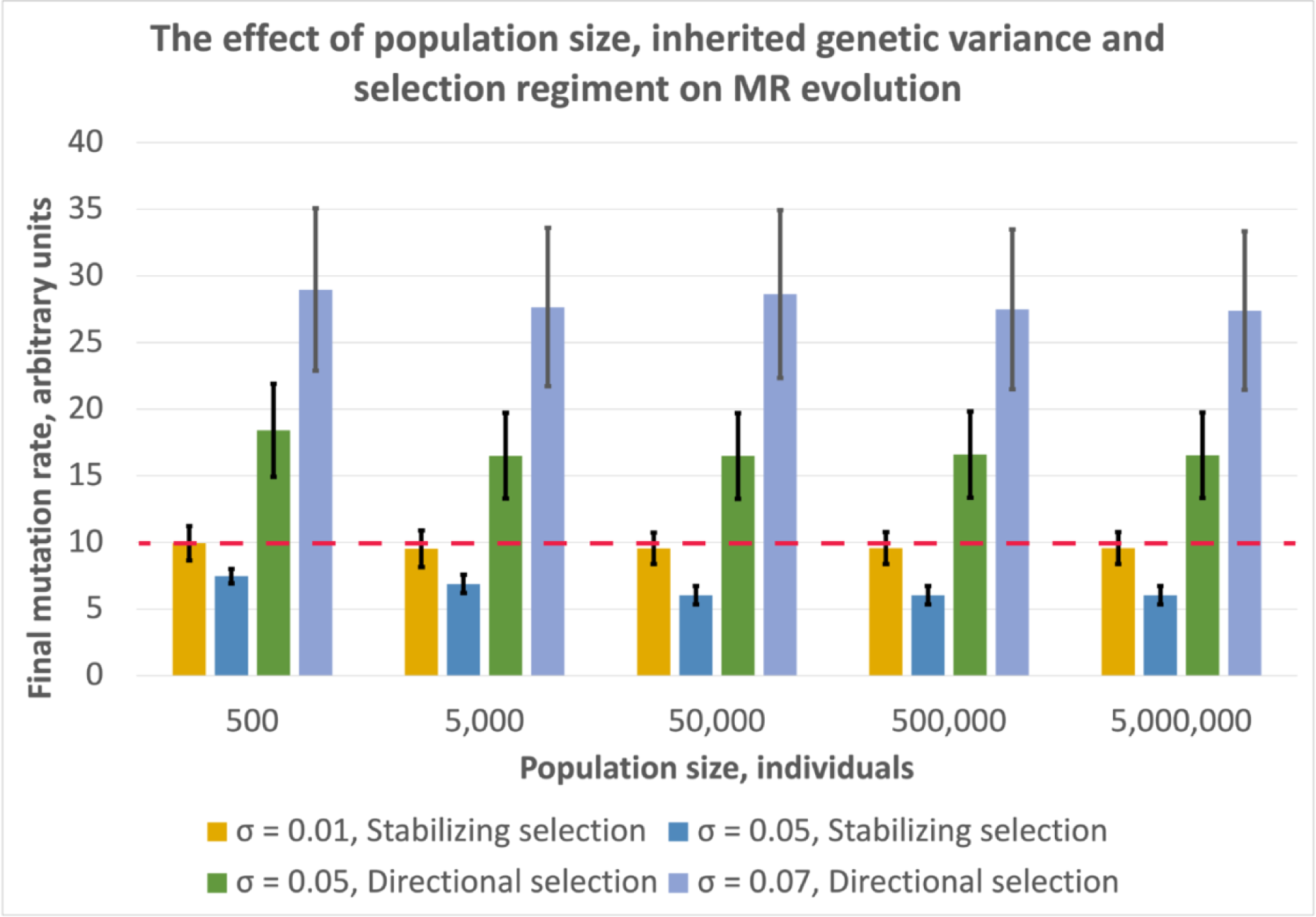
The effect of genetic drift on MR evolution. Inherited phenotypic variation is represented as the standard deviation of the distribution of phenotypic effects (DPhE) of mutations on the modeled adaptive quantitative trait and MR itself and is represented by the symbol ***σ***. Stabilizing and directional selection regimens acting on the modelled adaptive trait were applied with varying ***σ*** and population sizes under both selection conditions. The red dotted line indicates the initial MR at the beginning of simulations, which was 10 for all conditions.

In our simulated experiments (see **Fig. 8**), sample sizes are untypically large (∼500 through ∼5 million) and are also vastly different for within condition comparisons between different population sizes. This fact complicates most traditional statistical comparisons (sensitive to sample size) by rendering *p*-values practically meaningless. We did compare samples by applying both within- and between-condition pairwise comparisons using Student *t*- and related tests, resulting in practically all *p*-values being significant at alpha = 0.01. As to more descriptive statistics, due to the large sample sizes both standard errors of the mean (SEM) and even confidence intervals (CI) at alpha = 0.05 are uninformatively small and range in subdecimal numbers, with the latter being much smaller than the standard deviation (SD). We therefore plotted in **Fig. 8** the SD for error bars as a descriptive statistic illustrating the general variance and being among the least conservative statistics for the presented dataset. Therefore, it is interesting to notice that the data show almost total overlap of the range within one SD among all population sizes within each individual experimental selection regimen condition. On the other hand, there is absolutely no overlap within that range between selection regimen conditions. Given the very liberal nature of the chosen descriptive statistic and visual inspection of the chart in **Fig. 8**, the amount of genetic/phenotypic variation in a population is the main regulator of the amount of drift in the experimental population and also explains most of the variation in the simulated resulting MRs. Population size appears to have some contribution within the lower population size range of ∼500 to ∼5,000 individuals in most selection regimen conditions, but practically disappears into higher values. The latter observation could be explained by the fact that being roughly an inverse of the effective population size, the strength of *N_e_* induced drift follows hyperbolic decay as a function of population size and thus renders genetic drift a negligible force in populations with *N_e_* exceeding a few thousand individuals.

Beyond the main source of genetic drift that our results reveal, it is noteworthy that the regimen of selection acting on the simulated adaptive trait is the main determinant of the mode of selection acting on MR (see **Fig. 8**). In accord with Lynch’s overarching idea [1], the ability of stabilizing selection to lower MR is inversely related with the amount of genetic drift in a population (although drift appears to depend on standing variation more than on effective population size). The same principle appears to hold for directional selection as well. However, in support of our earlier findings (**Fig. 3**) and the main underlying hypothesis of this study, directional selection acting on population’s adaptive traits triggers an opposite mode of MR evolution compared to stabilizing. Variance in the resulting MR (shown with error bars in **Fig. 8**) appears to also behave in accord with this model, being highest under strong directional selection (coinciding with the highest MR) and lowest under strong stabilizing selection (coinciding with lowest MR).

## Discussion

Our results corroborate our initial hypothesis that the evolution of MRs in sexually reproducing populations is not universally directed toward lower MRs but is instead dependent on the general regimen of selection acting on adaptive traits. As shown in **Fig. 1**, stabilizing selection favoring adaptive trait expression closer to the population mean acts over generations to enrich the population with individuals harboring alleles that provide a lower MR, since such individuals tend to deviate less phenotypically from the favored phenotypic optimum. However, when directional selection is applied, favoring phenotypically deviant individuals from the tails of the phenotypic distribution, preferential survival is characteristic for individuals harboring alleles for higher MR, since such individuals tend to deviate in the expression of their adaptive trait further from the population mean. As a result (**Fig. 3**), we observe a reversal of the directionality of MR evolution. While the absolute magnitude of such a reversal will depend on many factors, such as the cost of higher MR, the amplitude of MR variation in a population or strength and efficiency of selection, our results still indicate that directional selection acting on adaptive traits at least counteracts the putative universal selection toward lower MRs. Evidence that selection regimen acting on adaptive traits has at least some role in modulating the evolution of MRs is further elucidated when we apply stronger directional selection and observe that its effect on MR evolution is further magnified (**Fig. 4A-B**).

Importantly, our results also indicate that MR evolution is regulated by a complex array of genetic factors rather than being primarily defined by a single population- or selection-related parameter. In particular, beyond the mode of selection acting on adaptive traits, our *in silico* experiments elucidate a tangible role for the cost of somatic mutations, multigenic inheritance of both MR and adaptive traits under selection, rates of genetic recombination, and genetic drift. Other factor may exist as well that we did not explore in this study. We expect, therefore, that because of the vast differences in biology, evolutionary status, and organization of genomes and populations among sexually reproducing taxa, different species should differ in which of these factors or combinations thereof are dominant in their taxon-specific process of MR evolution. Biological diversity is so colossal that a factor that would be common and overarching in MR evolution to all life might not even exist.

It is important to note, however, that as proposed by others [1,9–12], direct selection likely indeed always acts to lower MR because of the fitness cost of mutations. The cost of a higher MR should be manifested both in the increased phenotypic variance of offspring (with more offspring deviating from optimal trait values) and in the costs of concomitantly altered somatic MR, as germline and somatic MRs are linked, and mutations can contribute to aging, cancer and other fitness-limiting impairments [19,20,35–39]. Our previous modeling indicated that one mechanism to buffer the cost of somatic mutations is greater investment in somatic tissue maintenance, allowing for the evolution of higher MR and thus facilitating directional selection [40]. When we increased the fitness cost of mutations (**Fig. 4C-F**), it expectedly led to lower resulting MRs under both stabilizing and directional selection acting on the adaptive trait. The evolution of higher MRs in our study, based on the underlying hypothesis shown in **Fig. 1**, is not the result of direct selection. However, it is also not independent of selection. This process obviously results from the unique nature of MR as a phenotypic trait. Mutations negatively affect fitness of the current generation. However, MR is a trait that impacts the distributions/frequencies of most other traits, including adaptive traits that are under varying regimens of selection. Because of this effect, a certain form of co-selection for lower or higher MR appears to arise when major adaptive traits are under certain regimens of selection. Therefore, MR evolution appears always to be the result of two selective forces – the direct selection to lower MR because of the fitness cost of mutations and a second co-selective process imposed by selection regimen acting on adaptive traits and which acts regardless of MR fitness effects on the current generation. In the case of stabilizing selection, these two forces reinforce each other. Under directional selection they act in opposite directions, with the net result of this “compound selection” depending in each specific case on the relative strength of the two.

A substantial portion of phenotypic traits, including MR, are known to be encoded by multiple genes [41]. For MR, these include the myriad of genes involved in DNA replication and repair, as well as metabolic and stress response genes that modulate the chemical and biochemical intracellular environment (see Introduction). Our results reveal that the multigenic nature of both adaptive traits and MR has a profound effect on MR evolution. As we demonstrate in **Fig. 5**, multigenic inheritance of the adaptive trait buffers the ability of stabilizing selection to lower MR. This pattern can be explained by the lower number of mutations needed to alter a monogenic adaptive trait (just one gene to change), rendering a genotype more responsive to selection even with relatively lower MR. If the adaptive trait is multigenic (and we assume for simplicity roughly equal impact of each gene on the trait), deviation of such a trait from the selected population mean will require more mutations, and thus a higher MR is “tolerated” by stabilizing selection. Under directional selection, following the same logic, a multigenic adaptive trait leads to the evolution of a significantly higher MR compared to a monogenic trait, since more mutations are needed to alter a multigenic adaptive trait and therefore individuals that significantly deviate from the population mean phenotype (and favored by directional selection) are likely to harbor more alleles encoding a higher MR and/or alleles that have a more profound effect on MR. These results are also consistent with the mechanism proposed in **Fig. 1** and our initial hypothesis.

We further demonstrate (**Fig. 6**) that multigenic inheritance of MR also has a notable effect on MR evolution. Stabilizing selection is more efficient in lowering monogenically inherited MR. This pattern can be explained by the fact that selection does not directly act on genes, but rather it acts on phenotypic traits (a point often ignored since the early 1900s – but perforce the original Darwinian view). For monogenic traits, the encoding gene is directly visible to selection. While monogenic traits, such as mismatch repair deficiency caused by a single inactivating gene mutation (whether in yeast or humans), have been well described, it is likely that most variation in MR in a population cannot be ascribed to variation in a single gene [19,21,41]. Most population variability of MR in humans shows relatively moderate variation [21,22], although some extremes are indeed infrequently observed. There should exist strong selection against such monogenic mutations in natural populations. Modest changes in MR may confer a minimal cost – while selection may still be able to act on the overall MR phenotype, the strength of selection acting on alleles of any given gene contributing to MR (if highly multigenic) will be very weak or negligible. Together with co-selection for higher MR when selection favors deviant phenotypes at the tails of a phenotypic distribution, this reasoning can provide a solution for how higher MR can be co-selected despite its cost and given that evolution has “no eyes for the future”. In multigenic inheritance, the contribution of each single gene (assuming all genes have effects of comparable scale) to the resulting phenotype is smaller and can be compensated by other encoding genes. Therefore, stabilizing selection is less efficient at purging each individual unfavorable allele if MR is multigenic. We also observe (**Fig. 6C-D**) that directional selection acting on the adaptive trait promotes a higher MR when MR is multigenic. This can be explained following the same principle that the contribution of each individual allele encoding higher MR to the lower fitness of the resulting high-MR phenotype (due to the somatic and germline costs of mutations) is less visible to selection compared to MR encoded by a single gene. In fact, for monogenic encoded MR, a lower MR is obtained regardless of the selection regimen. Therefore high-MR alleles are harder for selection to purge from genotypes with multigenic MR encoding compared to monogenic. In addition, where more genes encode the adaptive trait and/or MR, recombination is less efficient at segregating these traits away from each other. These results further corroborate our initial hypothesis illustrated in **Fig. 1**.

As previously mentioned, genetic recombination is thought to prevent the evolution of higher MR in sexual species, since it introduces shuffling of genes in their position on chromosomes relative to other genes, thus effectively disrupting genetic linkage. Such shuffling prevents many genes from being co-selected such that one adaptive allele is favored by selection and others are passively co-selected by being physically located within the same genetic structure (such as for example a chromosome or a bacterial genome). Our results (**Fig. 7**) demonstrate indeed that the presence of genetic recombination buffers the ability of stabilizing selection to lower MR, as well as the ability of directional selection to promote the evolution of higher MRs, obviously because of the above-mentioned selection-independent randomization of the genetic location of alleles relative to each other. In essence, genetic recombination also acts as a factor affecting the balance of selection and genetic drift, because the process randomly shuffles pieces (at the gene level) of a whole phenotype that has survived (was selected), ensuring that the progeny will randomly differ from the (selected) parents. On the other hand, genetic recombination generates diversity which directly increases the overall efficiency of selection.

Interestingly, our results show a substantial portion of genetic drift (which also impacts MR evolution) comes from the population’s current genetic diversity (**Fig. 8**). The contribution of the effective population size (*N_e_*) to genetic drift appears to completely disappear in populations with the effective size exceeding a few thousand individuals and appears to be a tangible factor only in populations of hundreds and below. These results are consistent with the idea that while the effect of *N_e_* on the efficiency of selection is theoretically always present due to the sampling error phenomenon, its strength, being roughly an inverse of *N_e_*, rapidly deteriorates in the lower *N_e_* numbers by following a hyperbolic decay function of a linear increase in *N_e_*. On the other hand, the amount of genetic and the resulting phenotypic variation in a population is a factor directly regulating the efficiency of selection by altering the distribution and magnitude of selective coefficients in the population. Selective coefficients basically reflect phenotypic (and thus fitness) distances between genotypes/phenotypes under selection.

Roughly speaking, a population consisting of phenotypes that are exactly all the same will be completely dominated by drift, with zero efficiency of selection. Selection becomes more and more efficient as genetic/phenotypic diversity increases, because increased diversity means higher selective coefficients operating between specific phenotypes. The relationship between the two, shown in **Fig. 8**, appears to be more in a linear domain compared to that between genetic drift and *N_e_*. This relationship, together with the observed increases in MR under directional selection, implies that there should a *positive feedback loop* between MR and standing genetic variation during evolutionary transitions, whereby directional selection leads to the evolution of higher MR, higher MR then increases genetic variation in the population, which in turn increases the efficiency of selection and leads to even higher MRs and so on.

In line with this idea, there is evidence, for example, that MR has recently undergone a rapid evolution in humans [21]. Recent analyses of germline mutation rates across vertebrate taxa also surprisingly showed exceptionally high germline MRs in a number of domesticated animals (which have undergone recent periods of rapid directional selection) compared to wild species [42,43]. Notably, the negative correlation of MR with *N_e_*, consistent with the model proposed by Lynch [1] was reported to hold across taxa overall. Therefore, both the mechanism we propose and that proposed by Lynch [1] could be widespread in natural populations depending on selection regimens, with our prediction being that the effect of *N_e_* should increase during prolonged stabilizing selection periods acting on a population. On the other hand, our proposed ability of directional selection to indirectly select for higher MR could help explain the ability of very small populations, such as large tetrapods, to adapt to major environmental and rapid evolutionary transitions [44,45]. Some other examples include many species during colonization of novel environments when profound effects of genetic bottlenecks dramatically reduce a population’s genetic diversity and size (and therefore *N_e_*), but do not impair their ability to adapt rapidly, called the “genetic paradox of invasive species” [18,46–49]. Invasive species are thus a great example of a rapid adaptation in conditions of strong directional selection caused by altered environment and a concurrently reduced *N_e_*. Some studies provide evidence that evolution of elevated MR may contribute to success during biological invasions [18], which is consistent with our results. Adaptive change in the speciation process can also be quite rapid, often interrupting longer periods of relative stability of entire species [50]. Therefore, a number of theoretical frameworks have also been proposed earlier, such as “punctuated equilibria” [44,45] and Vrba’s cross-genealogical Turnover Pulse Hypothesis [51], or theoretical explorations of events of even larger-scale morphological change in evolution, such as Simpson’s Quantum Evolution [52] and Goldschmidt’s Macroevolution [53], to explain observed multiple events of rapid evolutionary transitions. Therefore, the mechanism we propose in this study could also be re-evaluated from the standpoint of other relevant theoretical frameworks. More recently, Lande [34,54] has considered the effects of population size, mutation, selection and other factors, on multigenic traits in peak shifts, allopatric speciation and evolutionary consequences of environmental change. Such formulations might well be re-evaluated in the light of our study. Perhaps, as well, a pathway for a better integration of the theoretical postulates of evo-devo with conventional selectionist evolutionary theory might also be enabled through a consideration of our results.

Overall, our study reveals a rather complex pattern of factors that likely impact MR evolution in sexually reproducing populations and raises a number of questions that warrant further research. For example, in real populations, many traits (like house-keeping physiological functions) are perhaps almost always under stabilizing selection, even when directional selection acts on some other trait. It is therefore unclear whether directional selection acting on one trait will be enough to promote the evolution of higher MRs when stabilizing selection acts on other traits and might buffer this effect depending on the relative cost of higher MR in relation to different traits in a complex phenotype. Intuitively, however, we can speculate that when one or some traits come under strong directional selection, the value of such traits is high from the standpoint of adaptation and survival and thus the negative effect of a higher MR on other (stabilized) traits might be better tolerated. Moreover, such scenarios could potentially trigger co-evolution of mechanisms buffering the effects of mutations on the compromised traits [55].

It is also unclear whether lifespan can be a factor in MR evolution [40], as perhaps the cost of somatic mutations might be lower for short-lived species, which could potentially amplify the process of higher MR evolution under directional selection regimens that we show here. There is recent evidence, for example, indicating that somatic MR scales inversely with lifespan across mammals [56].

And finally, it should be noted that our modeling is rather generic by nature and has some inherent limitations. For example, the effect of directional selection acting on monogenic non-quantitative traits on MR evolution will likely differ from the pattern we observed in this study, since those traits do not demonstrate a continuous distribution of phenotypic expression and instead selection works with their discrete frequencies. Also, our model only operates with a single generic species, and real parameter values of the sexual process will differ dramatically in different natural populations and evolutionary scenarios. The “species” we model does not represent any natural species. Therefore, we can expect, as already mentioned, that in certain specific cases of MR evolution the main underlying factors may differ from those we revealed in our study. Nonetheless, our study raises a number of fundamental novel questions in regard to MR evolution and the complexity of factors that underly it and should instigate more specific research, including experimental tests of the proposed processes.

## Acknowledgements

We thank John Thompson of the University of California Santa Cruz, Michael Lynch of Arizona State University and Maxim Zagoskin of the University of Tennessee for critical remarks and insightful suggestions on this work.

