## Supplements for "*In silico* experiments uncover a novel mechanism underlying mutation rate evolution in sexually reproducing populations"

**Supplement 1.**

***a) Model Matlab code***

```
clearvars -except

% STORAGE MATRICES
%=======================================
popDynStore = [];
averMutRate = [];
averAdaptTrait = [];
mutAdaptCorel = [];
highestFitnessPhenotype = [];
%=======================================

% INITIAL VARIABLES
%=======================================
maxTime = 10000;
popsize = 10000;
nMutGenes = 10;
nAdaptGenes = 10;
initAllele = 10;
initMutRate = 0.1;
inhVar = 0.05;
percentReproducing = 0.05;
recombinRate = 0.05;
envChange = 0.001;
targetPhenotype = initAllele;
targetChangeRate = initAllele*envChange;
tolerance = initAllele;
costOfMutRate = 0.001;
%=======================================

% MODE OF SELECTION
%=======================================
%targetPhenotypeGoal = targetPhenotype; % STABILIZING selection
targetPhenotypeGoal = Inf; % DIRECTIONAL selection
%=======================================

% INITIAL POPULATION
%=======================================
population = zeros(popsize, (nMutGenes*2) + (nAdaptGenes*2) + 1);
population(:,1) = 1 : popsize;
population(:,2 : nMutGenes*2+1) = normrnd(initAllele, inhVar*initAllele,
[popsize, nMutGenes*2]);
population(:, (nMutGenes*2+2) : (nMutGenes*2+2)+(nAdaptGenes*2-1)) = ...
    normrnd(initAllele, inhVar*initAllele, [popsize, nAdaptGenes*2]);
%========================================

% INITIAL CHROMOSOMES
%========================================
mutGenes1 = 2:1+nMutGenes;
mutGenes2 = 1+nMutGenes+1:1+nMutGenes+nMutGenes;
adaptGenes1 = 1+(nMutGenes*2)+1:1+(nMutGenes*2)+nAdaptGenes;
adaptGenes2 = 1+(nMutGenes*2)+nAdaptGenes+1:1+(nMutGenes*2)+(nAdaptGenes*2);
initMutRates = mean(population(:, [mutGenes1 mutGenes2]), 2);
```

```
initAdaptTraits = mean(population(:, [adaptGenes1 adaptGenes2]), 2);

% INITIAL DISTRIBUTION OF ADAPTIVE TRAIT AND MUTATION RATE
%========================================
initMutAdaptCorel = [initAdaptTraits(:,:).'; initMutRates(:,:).'];
%========================================



%--------------------------
%--------------------------
%----- SIMULATION RUN -----
%--------------------------
%--------------------------

for time = 1 : maxTime
    time

    % MUTATION
    %====================================
    mutMatrix = zeros(size(population, 1), size([mutGenes1 mutGenes2], 2));
    sigmas = mean(population(:, [mutGenes1 mutGenes2]),
2)./initAllele.*initMutRate;
    sigmas = repmat(sigmas, 1, size([mutGenes1 mutGenes2 adaptGenes1
adaptGenes2], 2));
    mutMatrix =...
        binornd(ones(size(population, 1), size([mutGenes1 mutGenes2
adaptGenes1 adaptGenes2], 2)), sigmas, size(population, 1),...
        size([mutGenes1 mutGenes2 adaptGenes1 adaptGenes2], 2));
    mutRates = mean(population(:, [mutGenes1 mutGenes2]), 2);
    mutRates = repmat(mutRates, 1, size([mutGenes1 mutGenes2 adaptGenes1
adaptGenes2], 2));
    mutVariance = inhVar*mutRates;
    mutVariance = mutVariance.*mutMatrix;
    population(:, [mutGenes1 mutGenes2 adaptGenes1 adaptGenes2]) = ...
        population(:, [mutGenes1 mutGenes2 adaptGenes1 adaptGenes2]) +...
        normrnd(0, mutVariance, size(mutVariance, 1), size(mutVariance, 2));
    population(population < 0) = 0;
    %===================================



    % RECOMBINATION
    %===================================

        % mut rate alleles recombine
    recMatrixMut = binornd(1, recombinRate, [size(population, 1),
nMutGenes]);
    recMatrixMut(recMatrixMut > 1) = 1;
    backCopyMut1 = population(:, mutGenes1);
    backCopyMut2old = population(:, mutGenes2);
    backCopyMut2 = population(:, mutGenes2);
    backCopyMut2(recMatrixMut(:, :)==1) = backCopyMut1(recMatrixMut(:,
:)==1);
    backCopyMut1(recMatrixMut(:, :)==1) = backCopyMut2old(recMatrixMut(:,
:)==1);
    population(:, mutGenes1) = backCopyMut1;
```

```
    population(:, mutGenes2) = backCopyMut2;

        %adaptive alleles recombine
    recMatrixAdapt = binornd(1, recombinRate, [size(population, 1),
nAdaptGenes]);
    recMatrixAdapt(recMatrixAdapt > 1) = 1;
    backCopyAdapt1 = population(:, adaptGenes1);
    backCopyAdapt2old = population(:, adaptGenes2);
    backCopyAdapt2 = population(:, adaptGenes2);
    backCopyAdapt2(recMatrixAdapt(:, :)==1) =
backCopyAdapt1(recMatrixAdapt(:, :)==1);
    backCopyAdapt1(recMatrixAdapt(:, :)==1) =
backCopyAdapt2old(recMatrixAdapt(:, :)==1);
    population(:, adaptGenes1) = backCopyAdapt1;
    population(:, adaptGenes2) = backCopyAdapt2;
    %===================================



    % REPRODUCTION
    %===================================

        % population is shuffled and sorted to take random individuals for
reproduction
    population(:, 1) = randperm(size(population, 1), size(population, 1)).';
    population = sortrows(population);

        % number of reproducing individuals determined
    if(mod(floor(size(population, 1)*percentReproducing), 2) == 0)
        reprodPopNum = floor(size(population, 1)*percentReproducing);
    else
        reprodPopNum = floor(size(population, 1)*percentReproducing)-1;
    end

    if(reprodPopNum > 0)

        progeny = zeros(reprodPopNum/2, size(population, 2));
        progeny(:,1) = [1:size(progeny, 1)];

        % gamete segregation
        chromosomeSegregation = binornd(1, 0.5, [1, reprodPopNum]);
        for i = 1 : 2 : reprodPopNum
            if(chromosomeSegregation(i)==1)
                progeny((i+1)/2, mutGenes1) = population(i, mutGenes1);
                progeny((i+1)/2, adaptGenes1) = population(i, adaptGenes1);
            else
                progeny((i+1)/2, mutGenes1) = population(i, mutGenes2);
                progeny((i+1)/2, adaptGenes1) = population(i, adaptGenes2);
            end

            if(chromosomeSegregation(i+1)==1)
                progeny((i+1)/2, mutGenes2) = population(i+1, mutGenes1);
                progeny((i+1)/2, adaptGenes2) = population(i+1, adaptGenes1);
            else
                progeny((i+1)/2, mutGenes2) = population(i+1, mutGenes2);
                progeny((i+1)/2, adaptGenes2) = population(i+1, adaptGenes2);
```

```
            end
        end

        % progeny added to the population
        population = [population; progeny];
        population(:, 1) = [1:size(population, 1)];
    end
    %===================================



    % SURVIVAL
    %===================================

        % determining relative individual fitness
    mutRates = mean(population(:, [mutGenes1 mutGenes2]), 2);
    adaptTraits = mean(population(:, [adaptGenes1 adaptGenes2]), 2);
    distances = abs(adaptTraits(:)-targetPhenotype);
    mutFactors = 1 + (((mutRates(:)./initAllele) - 1).*costOfMutRate);
    distances = distances.*mutFactors;
    fitness = (1 - (distances(:)./tolerance));
    fitness(fitness < 0) = 0;

        % mortality
    overkill = size(population, 1)/popsize;
    if(overkill > 1)
        fitness = fitness.*(1/overkill);
        fitness = 1 - fitness;
        capdeath = binornd(1, fitness);
        population(capdeath == 1, :) = 0;
        population = population(population(:, 1) > 0, :);
    end
    %=====================================



    % UPDATES
    %=====================================

        % highest fitness phenotype updated
    if(targetPhenotype < targetPhenotypeGoal)
        targetPhenotype = targetPhenotype + targetChangeRate;
    end
    highestFitnessPhenotype(time) = targetPhenotype;

        % current population data are recorded
    popDynStore(time) = size(population, 1);
    averMutRate(time) = round(real(mean(mutRates(:, 1)))*1000)/1000;
    averAdaptTrait(time) = round(real(mean(adaptTraits(:, 1)))*1000)/1000;
    x=randi(size(population, 1), [1, 100]);
    mutAdaptCorel = [mutAdaptCorel, [adaptTraits(x,:).'; mutRates(x,:).']];
    %=====================================
end
```

**Supplement 2**

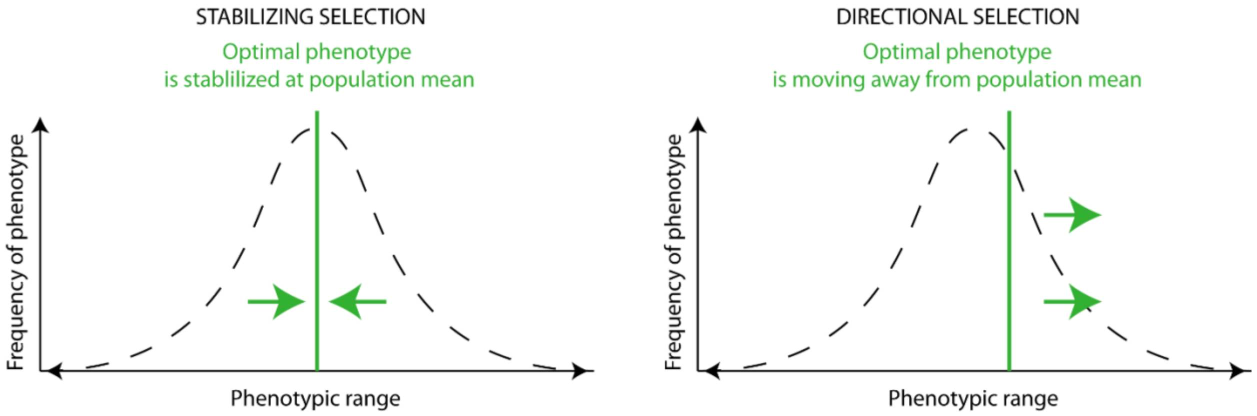


**Mode of selection**. Two modes of selection are modeled: stabilizing selection (left chart) and directional selection (right chart). Under stabilizing selection, the initial population mean phenotype is the highest fitness phenotype (green vertical line) and is kept fixed throughout the entire simulation run. Under directional selection, the highest fitness phenotype is moving over time (with each simulation step) away from the initial population mean and toward the selected phenotypic tail. When selective elimination of excess population is applied, phenotypes that are closer to the current optimal phenotype have higher chances of survival in each trial (each simulation step).

**Supplement 3a to Fig. 3A-B**

*Parameters*:

| | |
|---|---|
| Population size | 10000 |
| Number of mutation rate genes | 10 |
| Number of adaptive trait genes | 10 |
| Inherited variance | 0.05 |
| Recombination rate | 0.05 |
| Environmental change rate | 0.0013 |
| Mode of selection (stabilizing vs directional) | directional |
| Cost of mutation rate | 0.001 |

| Directional selection on adaptive trait | Stronger directional selection on adaptive trait |
|---|---|

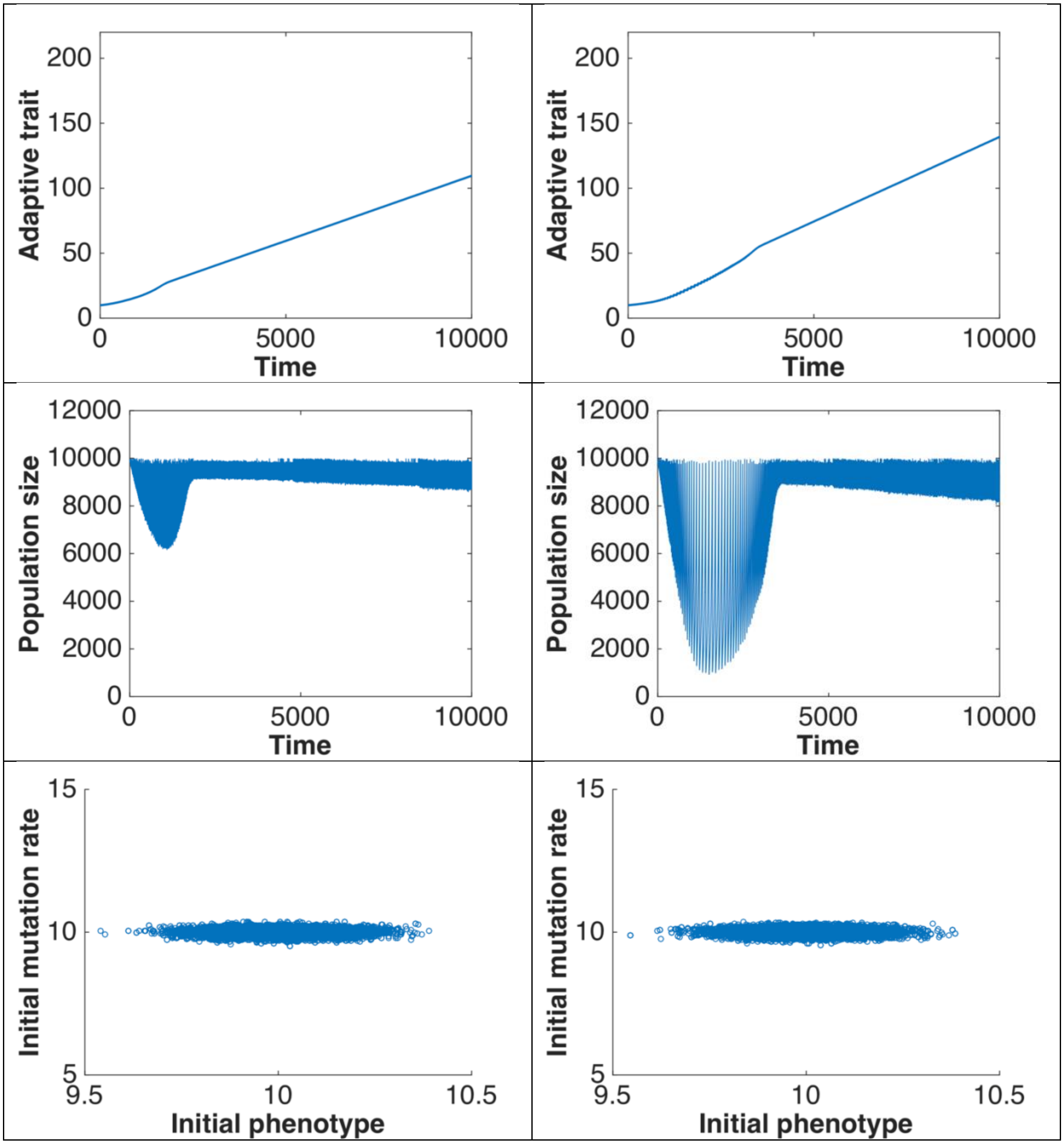
Adaptive trait
Time
0
50
100
150
200
5000
10000
Adaptive trait
Time
0
50
100
150
200
5000
10000
Population size
Time
0
2000
4000
6000
8000
10000
12000
5000
10000
Population size
Time
0
2000
4000
6000
8000
10000
12000
5000
10000
Initial mutation rate
Initial phenotype
5
10
15
9.5
10
10.5
Initial mutation rate
Initial phenotype
5
10
15
9.5
10
10.5

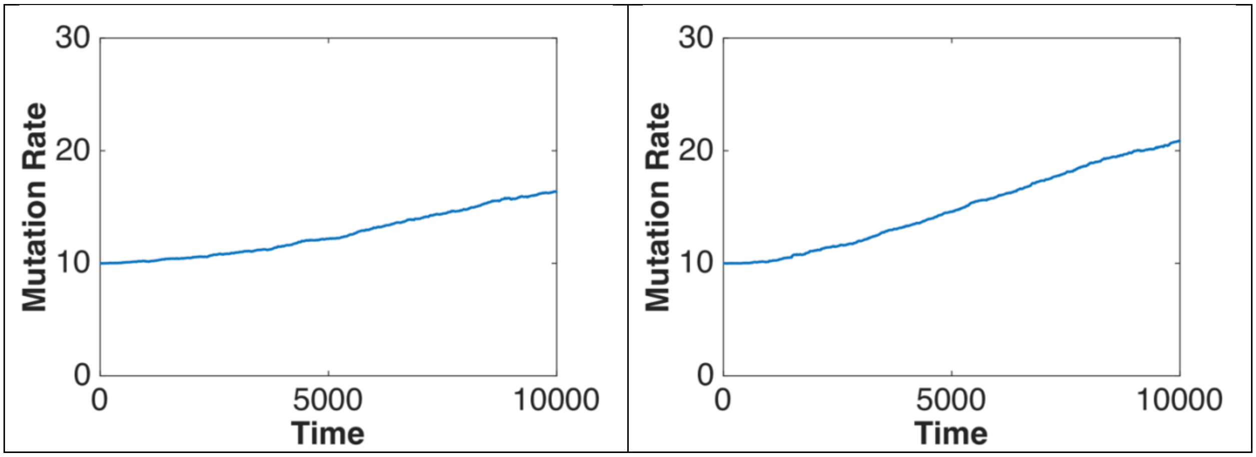

30
20
10
0
Mutation Rate
0
5000
10000
Time
30
20
10
0
Mutation Rate
0
5000
10000
Time

**Supplement 3b to Fig. 3C-D**

*Parameters*:

| | |
|---|---|
| Population size | 10000 |
| Number of mutation rate genes | 10 |
| Number of adaptive trait genes | 10 |
| Inherited variance | 0.05 |
| Recombination rate | 0.05 |
| Environmental change rate | 0.001 |
| Mode of selection (stabilizing vs directional) | stabilizing |
| Cost of mutation rate | 0.1 |

| Stabilizing selection on adaptive trait | Stabilizing selection on adaptive trait at higher cost of mutations |
|---|---|
| 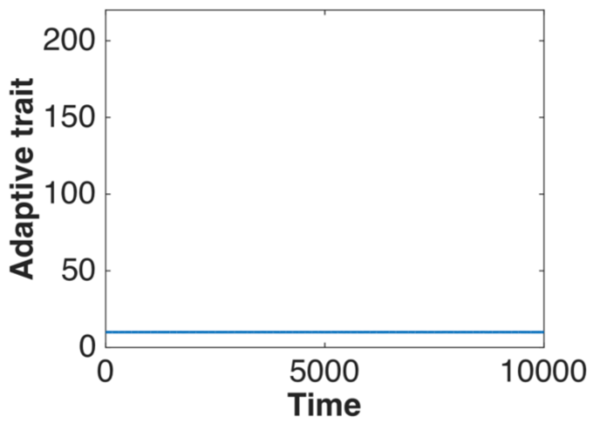 | 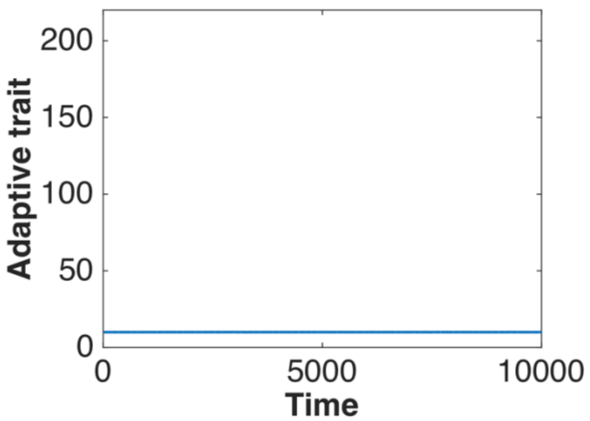 |
| 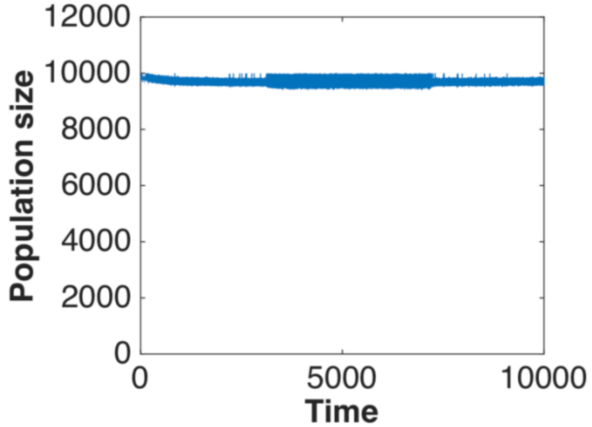 | 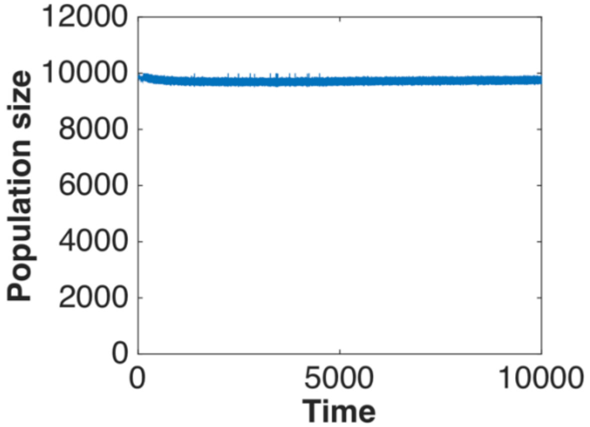 |

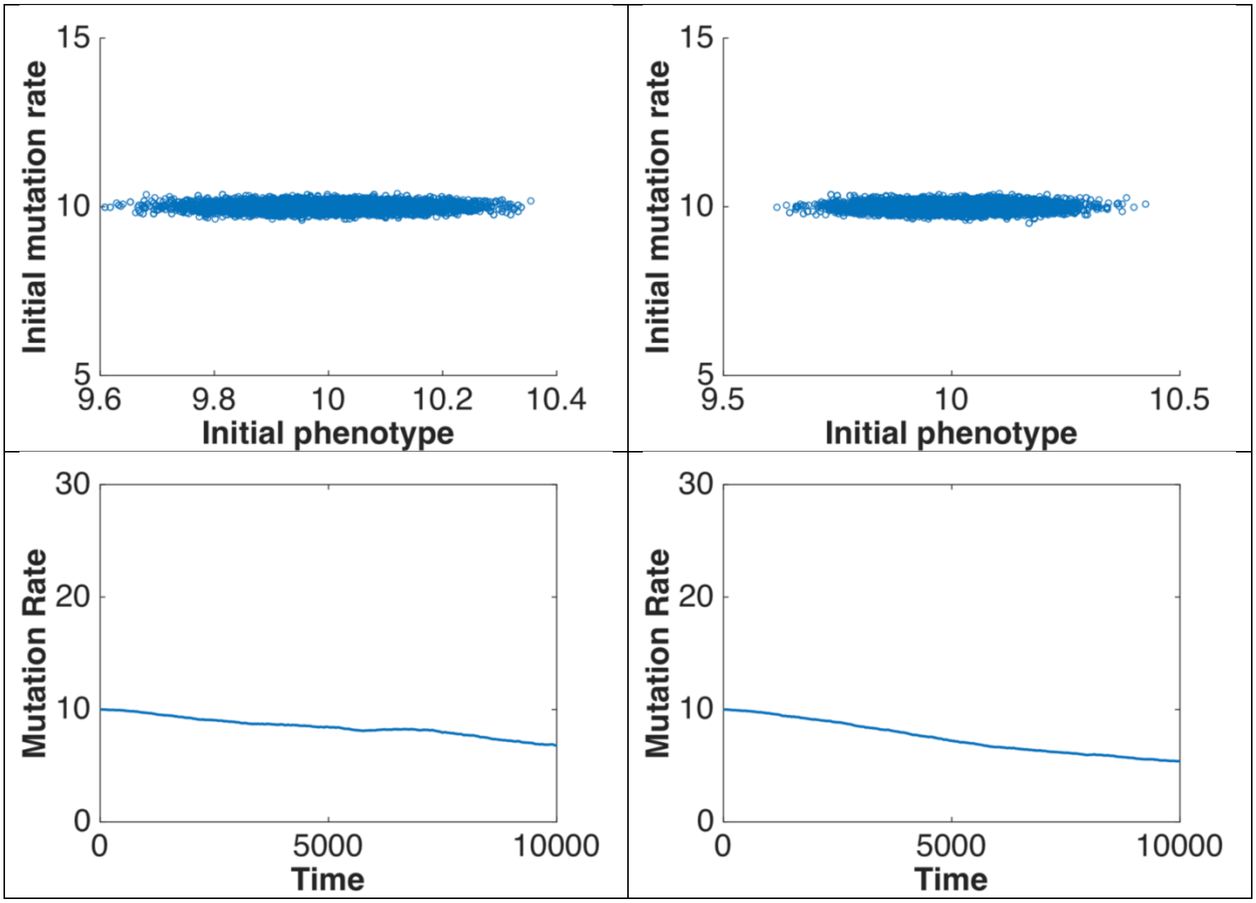

15
10
5
Initial mutation rate
9.6
9.8
10
10.2
10.4
Initial phenotype
15
10
5
Initial mutation rate
9.5
10
10.5
Initial phenotype
30
20
10
0
Mutation Rate
0
5000
10000
Time
30
20
10
0
Mutation Rate
0
5000
10000
Time

**Supplement 3c to Fig. 3E-F**

*Parameters*:

| | |
|---|---|
| Population size | 10000 |
| Number of mutation rate genes | 10 |
| Number of adaptive trait genes | 10 |
| Inherited variance | 0.05 |
| Recombination rate | 0.05 |
| Environmental change rate | 0.001 |
| Mode of selection (stabilizing vs directional) | directional |
| Cost of mutation rate | 0.1 |

| Directional selection on adaptive trait | Directional selection on adaptive trait at higher cost of mutations |
|---|---|
| 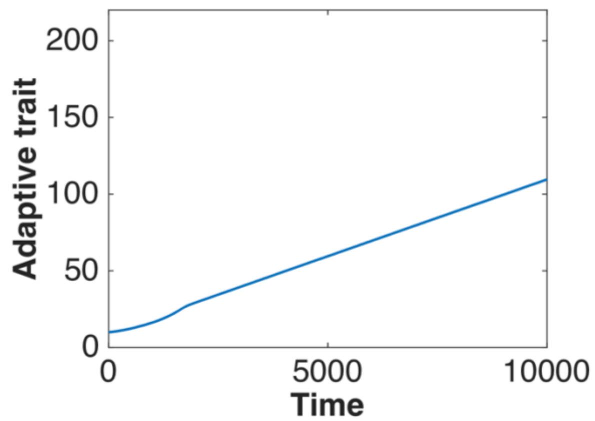  | 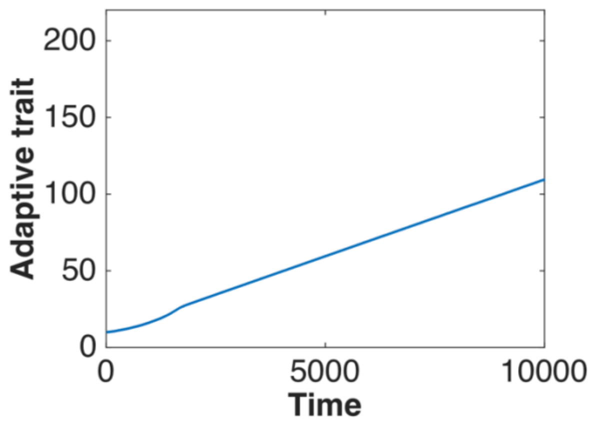  |
| 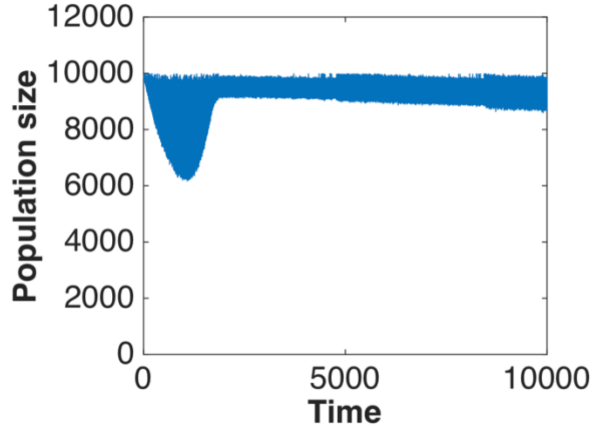  | 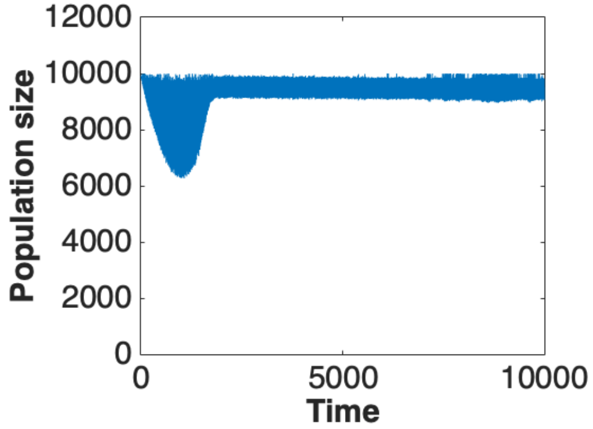  |

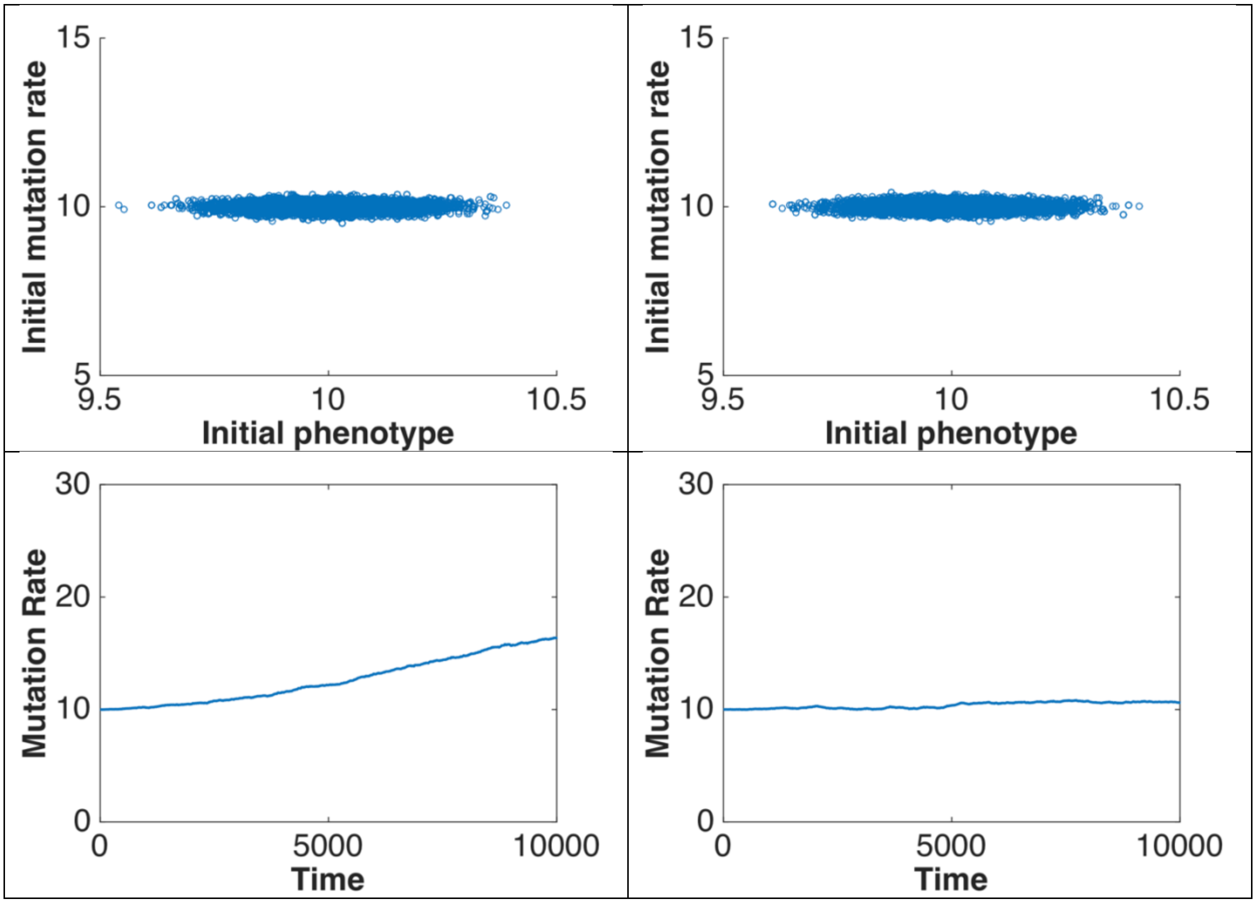

15
10
5
Initial mutation rate
9.5
10
10.5
Initial phenotype
15
10
5
Initial mutation rate
9.5
10
10.5
Initial phenotype
30
20
10
0
Mutation Rate
0
5000
10000
Time
30
20
10
0
Mutation Rate
0
5000
10000
Time

**Supplement 4a to Fig. 4A-B**

*Parameters*:

| | |
|---|---|
| Population size | 10000 |
| Number of mutation rate genes | 10 |
| Number of adaptive trait genes | 1 |
| Inherited variance | 0.05 |
| Recombination rate | 0.05 |
| Environmental change rate | 0.001 |
| Mode of selection (stabilizing vs directional) | stabilizing |
| Cost of mutation rate | 0.001 |

| Stabilizing selection on adaptive trait) | Stabilizing selection on a monogenic adaptive trait |
|---|---|
| 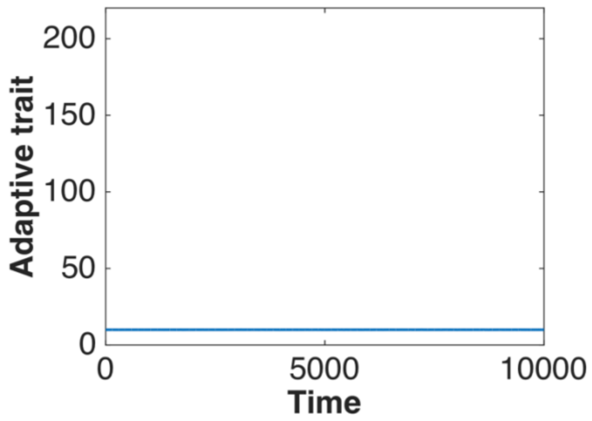 | 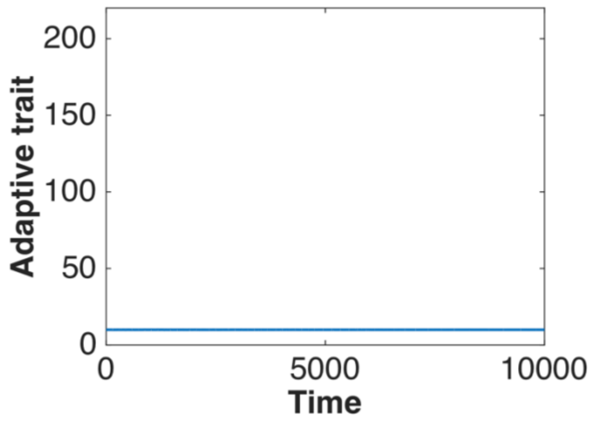 |
| 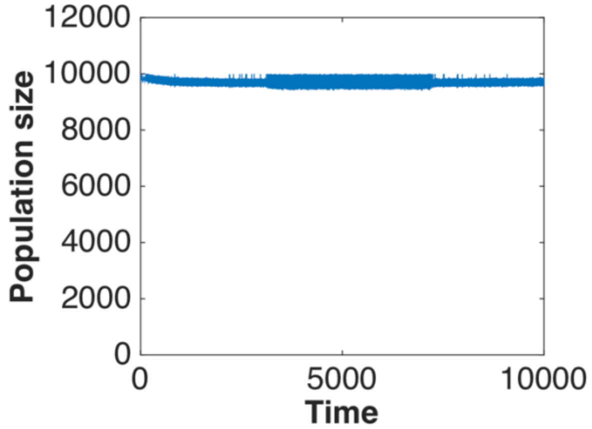 | 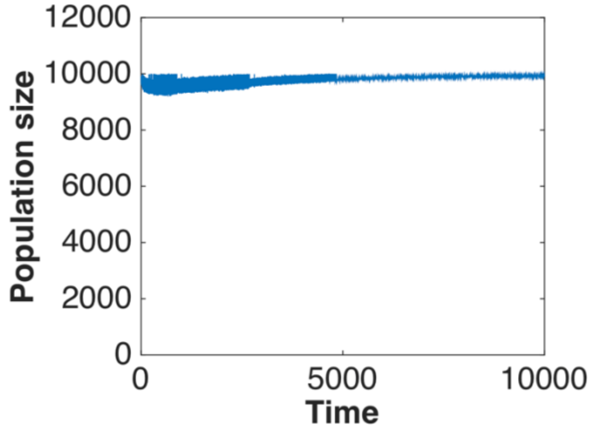 |

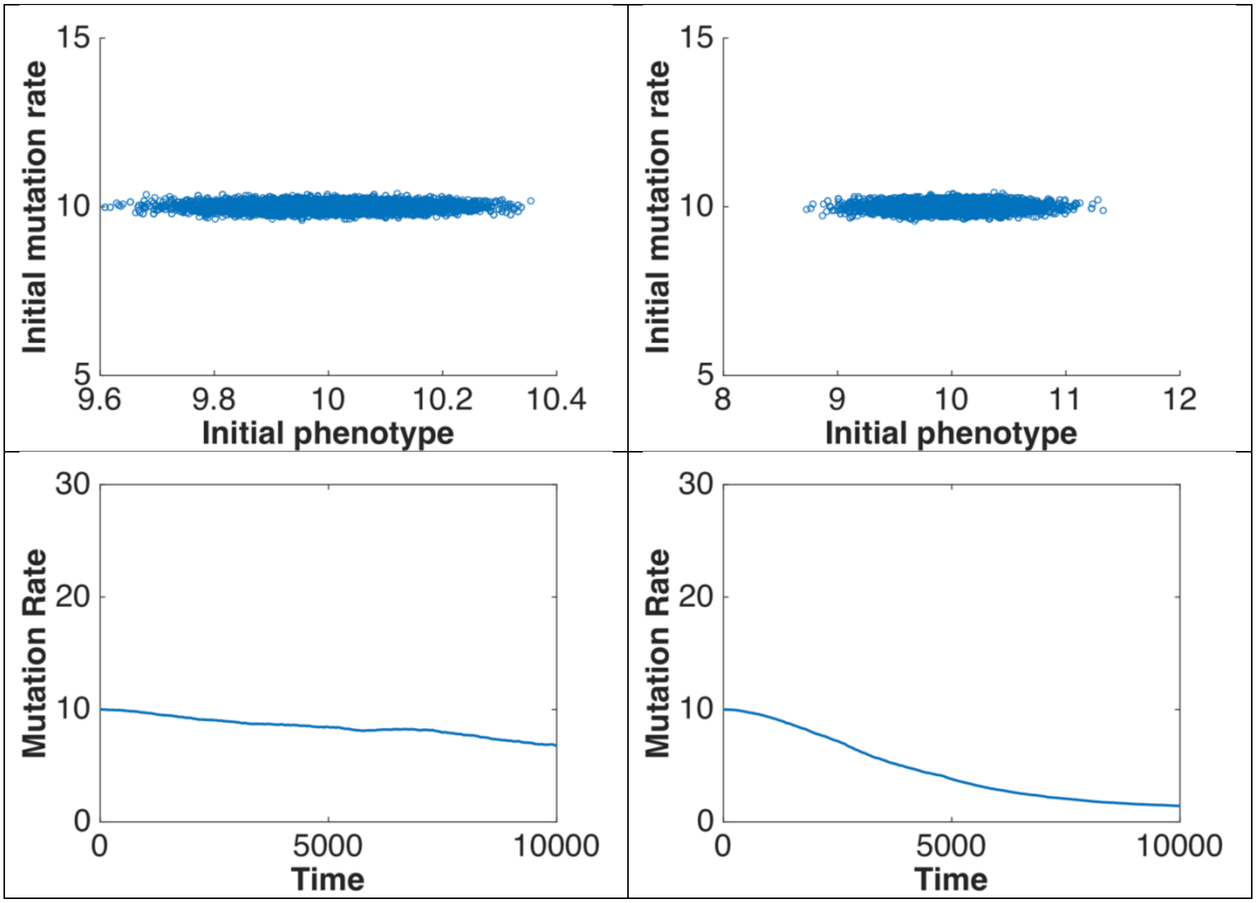

Initial mutation rate
15
10
5
9.6
9.8
10
10.2
10.4
Initial phenotype
Initial mutation rate
15
10
5
8
9
10
11
12
Initial phenotype
Mutation Rate
30
20
10
0
0
5000
10000
Time
Mutation Rate
30
20
10
0
0
5000
10000
Time

**Supplement 4b to Fig. 4D-E**

*Parameters:*

| | |
|---|---|
| Population size | 10000 |
| Number of mutation rate genes | 10 |
| Number of adaptive trait genes | 1 |
| Inherited variance | 0.05 |
| Recombination rate | 0.05 |
| Environmental change rate | 0.001 |
| Mode of selection (stabilizing vs directional) | directional |
| Cost of mutation rate | 0.001 |

| Directional selection on adaptive trait | Directional selection on a monogenic adaptive trait |
|---|---|

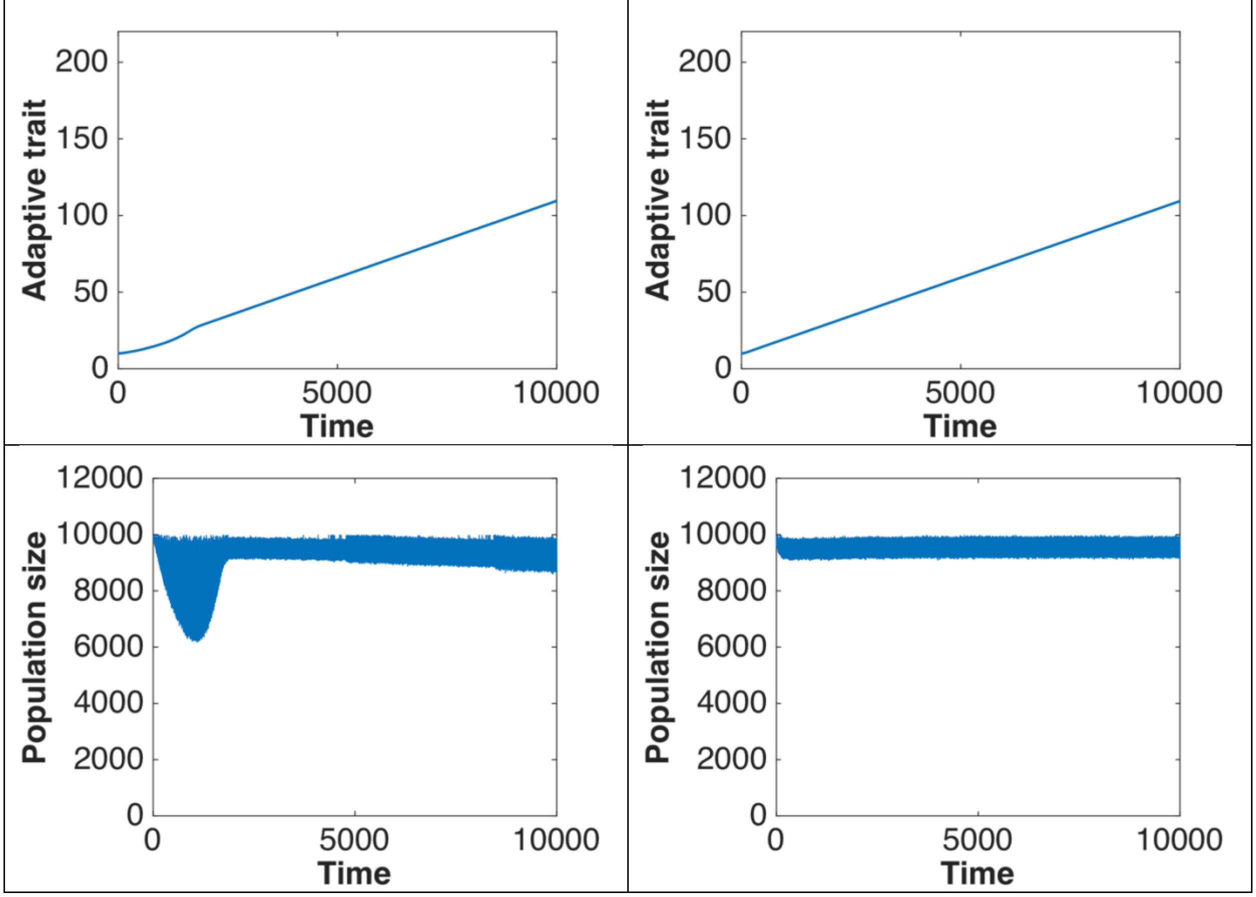

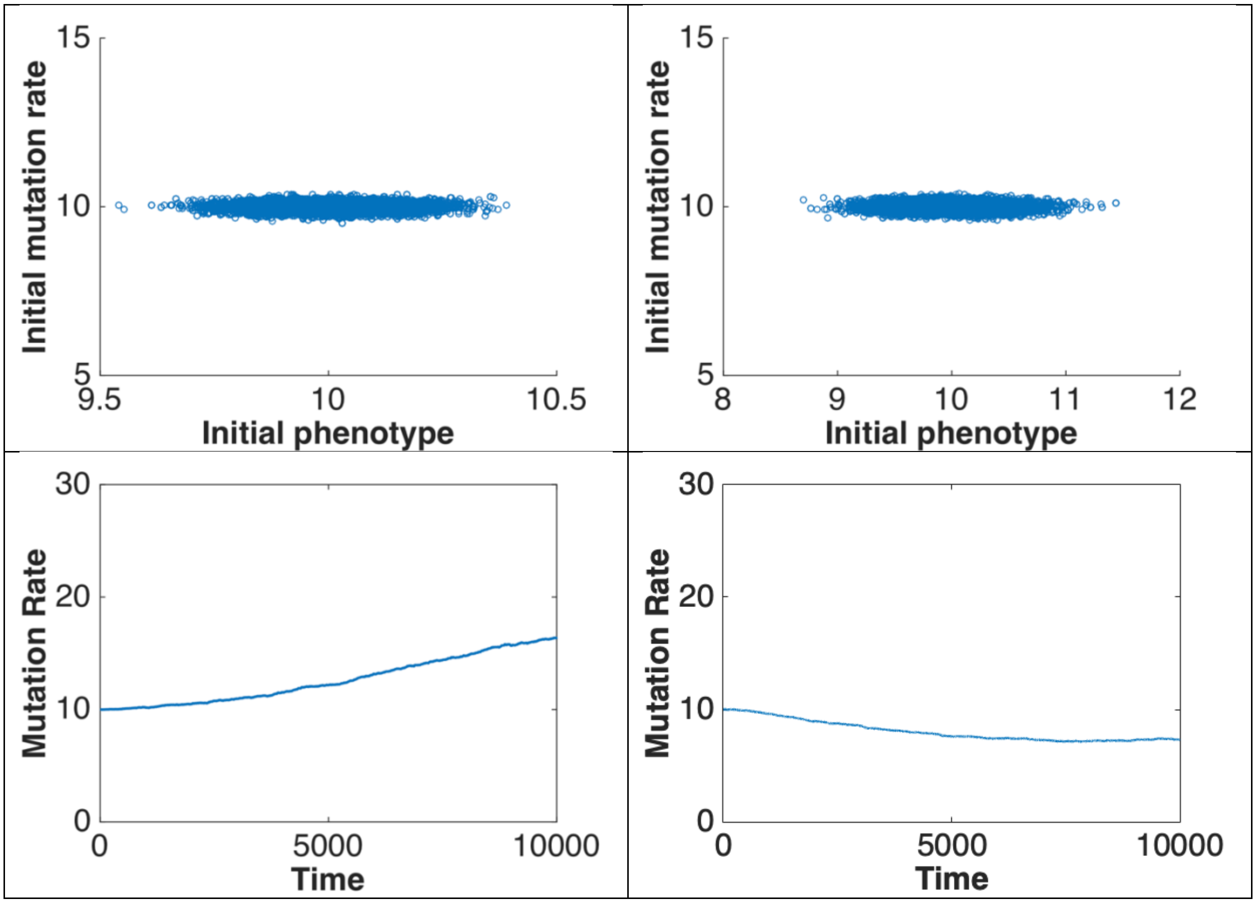

15
10
5
9.5
10
10.5
Initial mutation rate
Initial phenotype
15
10
5
8
9
10
11
12
Initial mutation rate
Initial phenotype
30
20
10
0
0
5000
10000
Mutation Rate
Time
30
20
10
0
0
5000
10000
Mutation Rate
Time

**Supplement 4c to Fig. 4C**

*Parameters*:

| | |
|---|---|
| Population size | 10000 |
| Number of mutation rate genes | 10 |
| Number of adaptive trait genes | 1 |
| Inherited variance | 0.01 |
| Recombination rate | 0.05 |
| Environmental change rate | 0.001 |
| Mode of selection (stabilizing vs directional) | stabilizing |
| Cost of mutation rate | 0.001 |

| Stabilizing selection on a monogenic adaptive trait | Stabilizing selection on a monogenic adaptive trait at narrow distribution of mutation phenotypic effects |
|---|---|
| 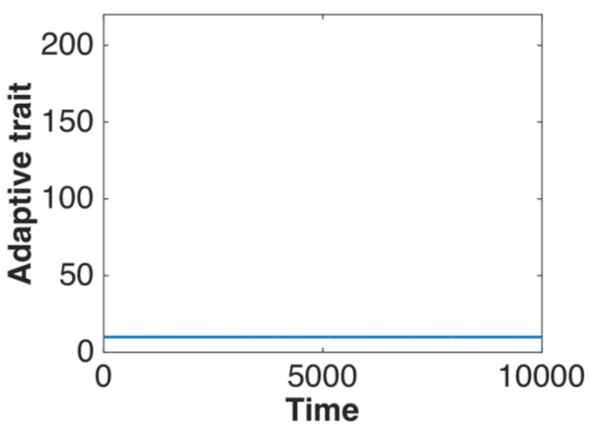 | 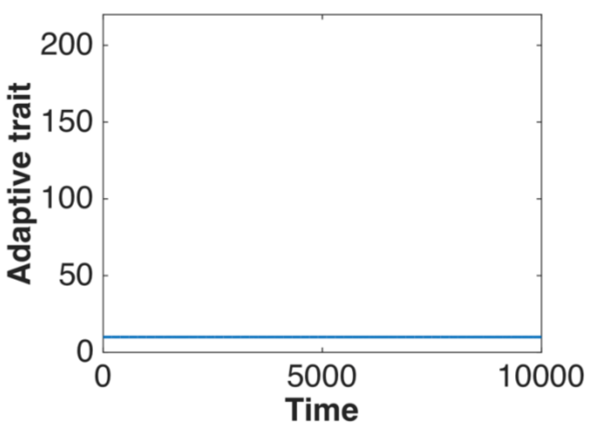 |
| 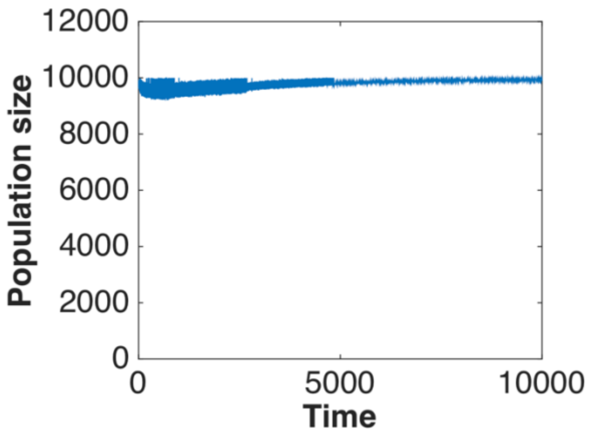 | 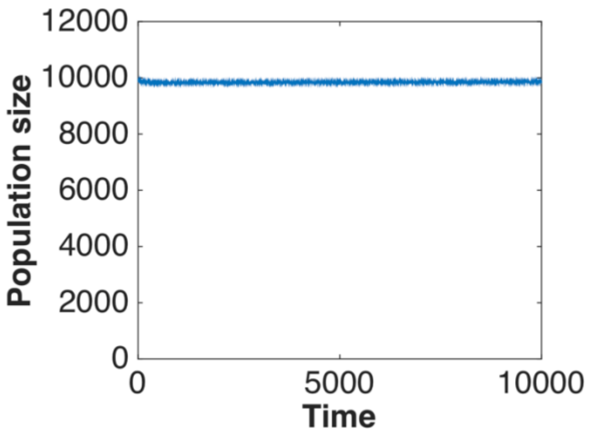 |

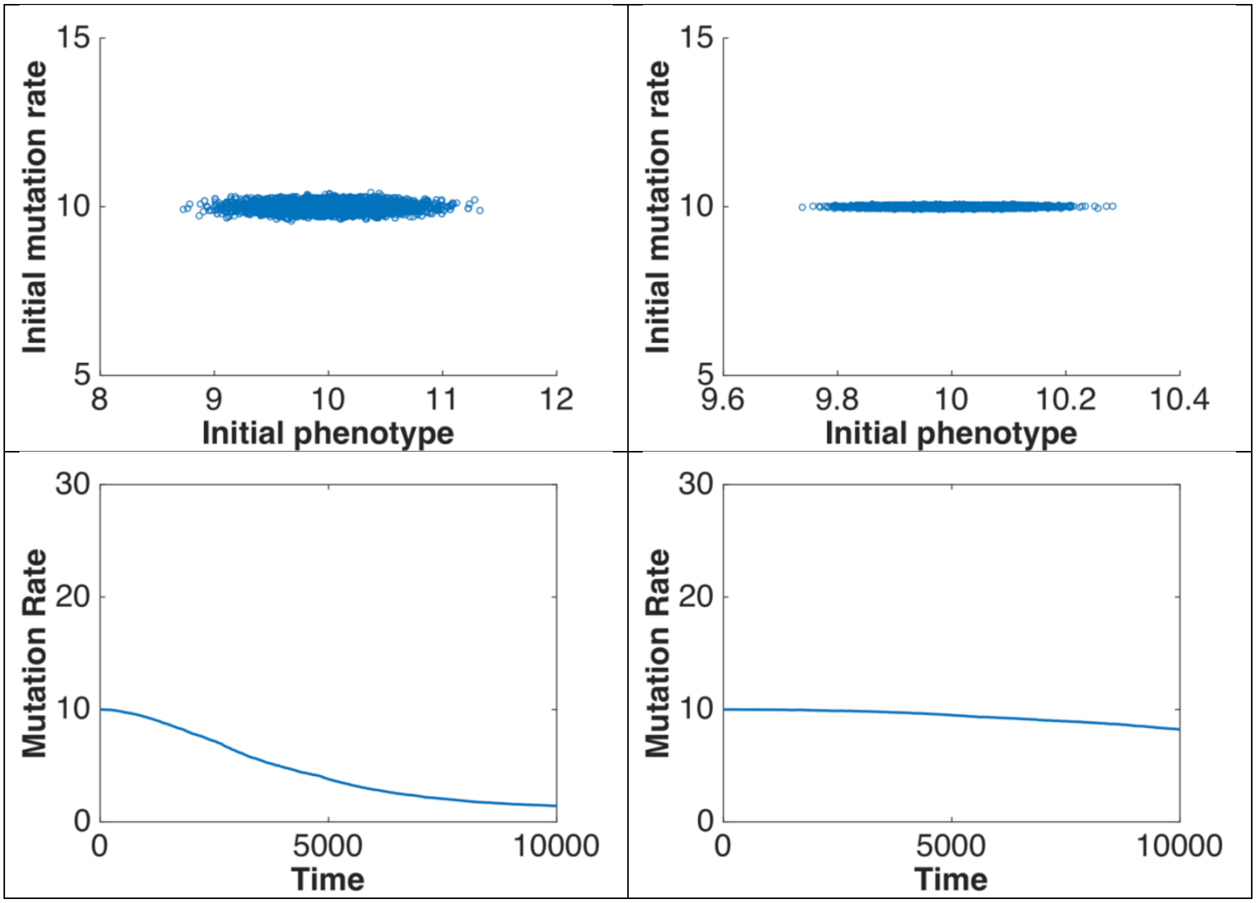

Initial mutation rate
15
10
5
8
9
10
11
12
Initial phenotype
Initial mutation rate
15
10
5
9.6
9.8
10
10.2
10.4
Initial phenotype
Mutation Rate
30
20
10
0
0
5000
10000
Time
Mutation Rate
30
20
10
0
0
5000
10000
Time

**Supplement 4d to Fig. 4F**

*Parameters*:

| | |
|---|---|
| Population size | 10000 |
| Number of mutation rate genes | 10 |
| Number of adaptive trait genes | 1 |
| Inherited variance | 0.018 |
| Recombination rate | 0.05 |
| Environmental change rate | 0.001 |
| Mode of selection (stabilizing vs directional) | directional |
| Cost of mutation rate | 0.001 |

| Directional selection on a monogenic adaptive trait | Directional selection on a monogenic adaptive trait at narrow distribution of mutation phenotypic effects |
|---|---|
| 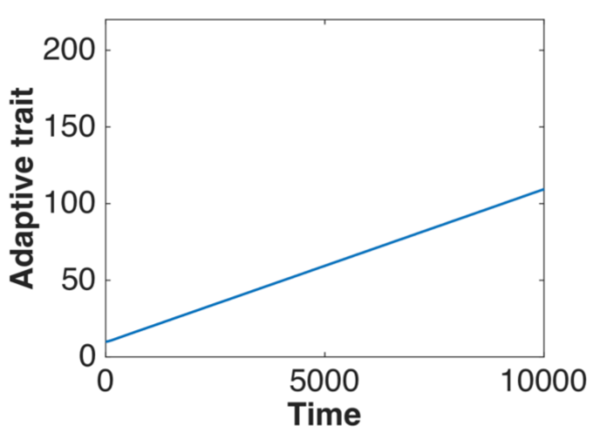  | 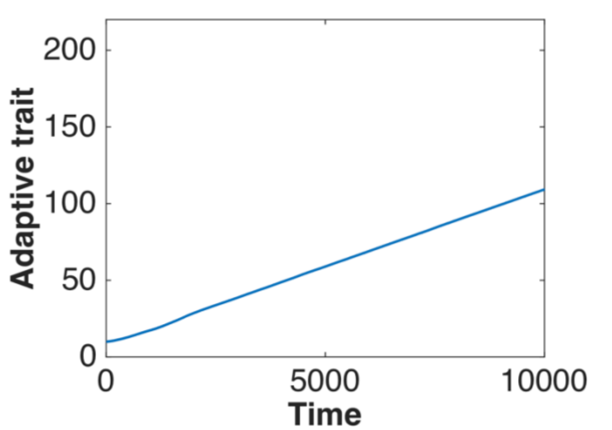  |
| 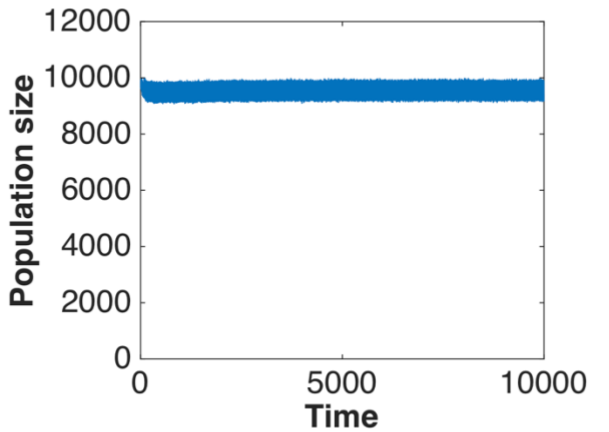  | 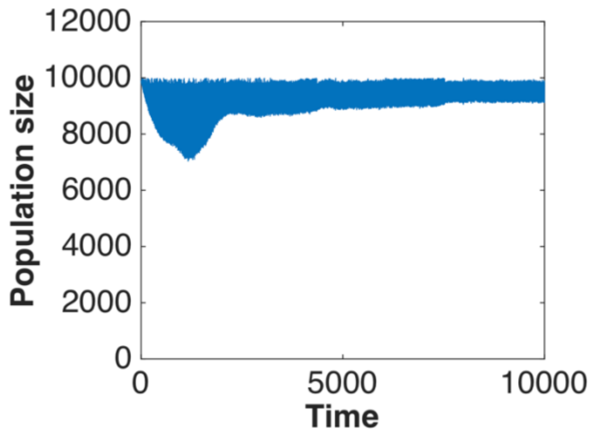  |

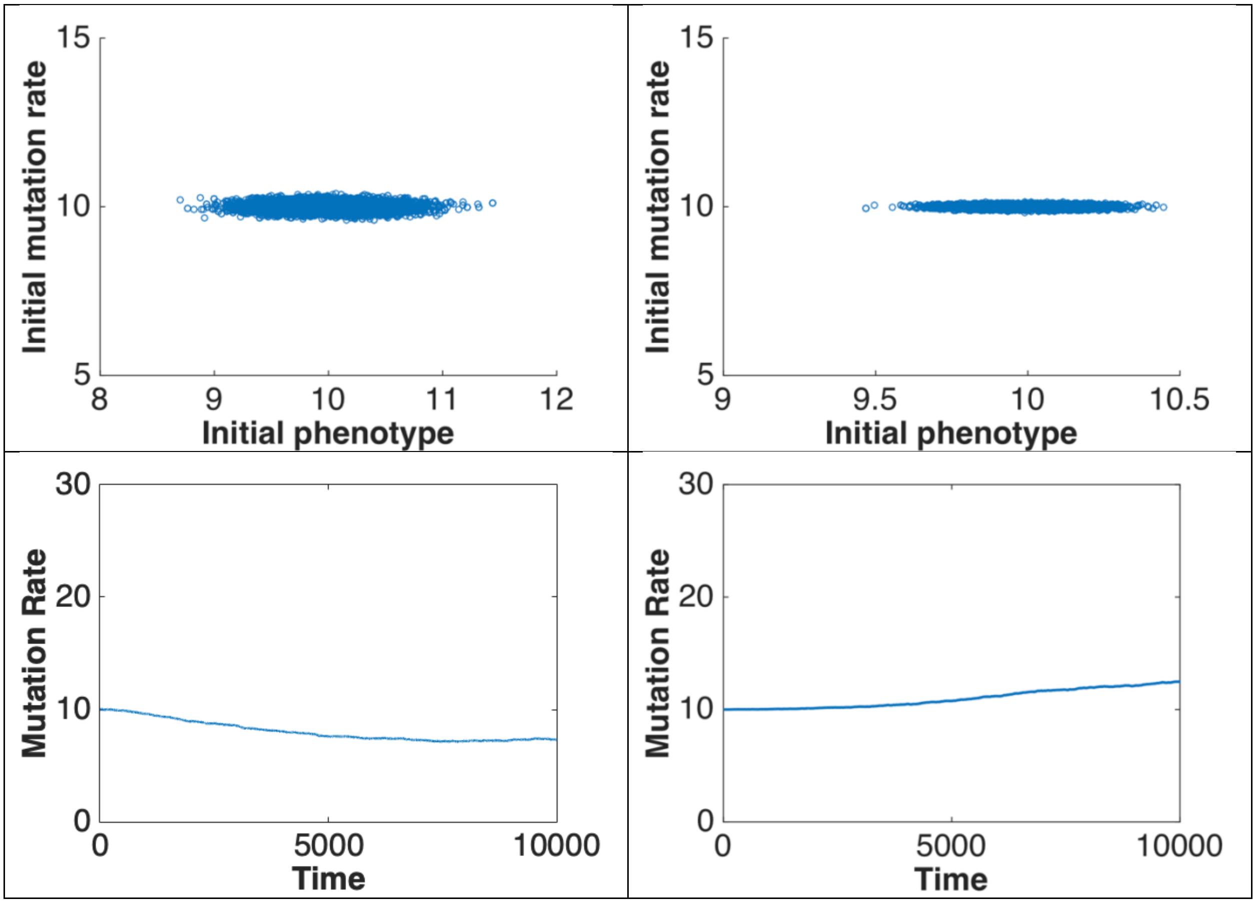

Initial mutation rate
Initial phenotype
15
10
5
8
9
10
11
12
Initial mutation rate
Initial phenotype
15
10
5
9
9.5
10
10.5
Mutation Rate
Time
30
20
10
0
0
5000
10000
Mutation Rate
Time
30
20
10
0
0
5000
10000

**Supplement 5a to Fig. 5A-B**

*Parameters:*

| | |
|---|---|
| Population size | 10000 |
| Number of mutation rate genes | 1 |
| Number of adaptive trait genes | 10 |
| Inherited variance | 0.05 |
| Recombination rate | 0.05 |
| Environmental change rate | 0.001 |
| Mode of selection (stabilizing vs directional) | stabilizing |
| Cost of mutation rate | 0.001 |

| Stabilizing selection on adaptive trait | Stabilizing selection on adaptive trait at monogenic inheritance of mutation rate |
|---|---|
| 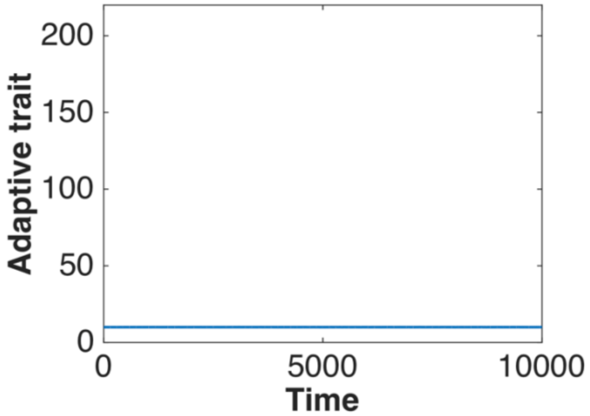 | 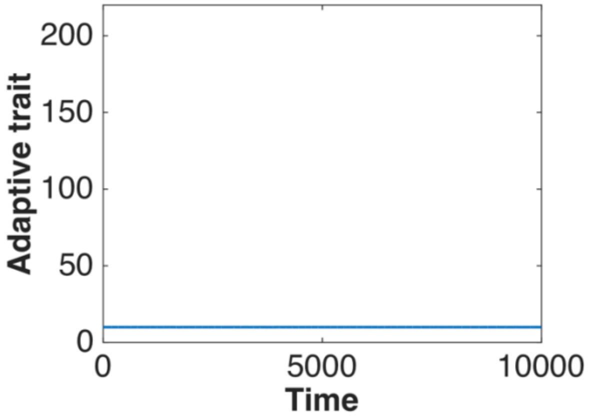 |
| 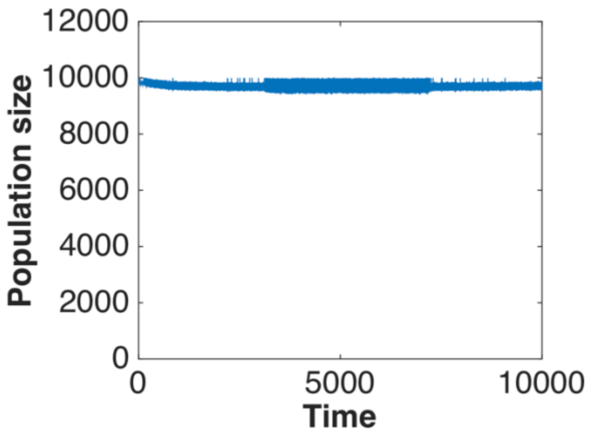 | 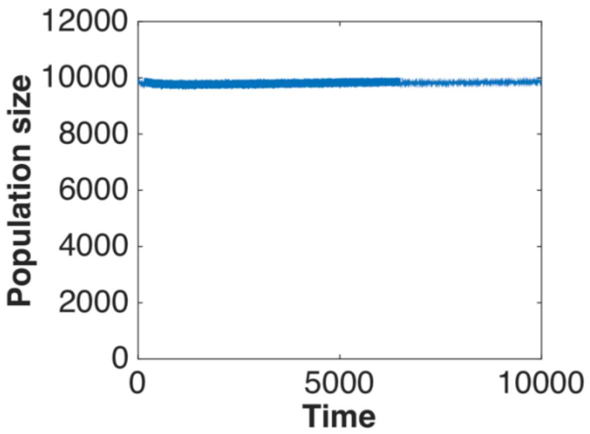 |

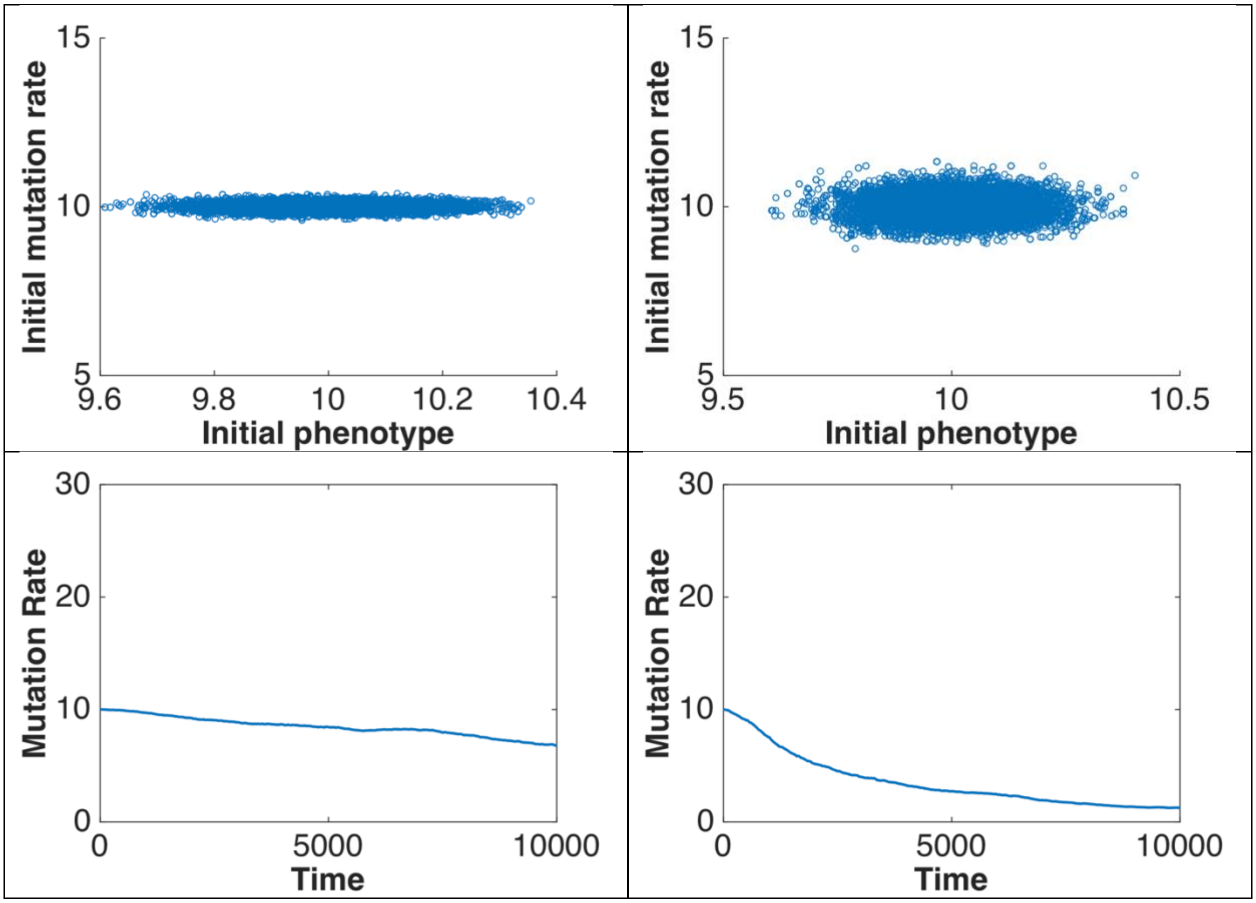

Initial mutation rate
Initial phenotype
Initial mutation rate
Initial phenotype
Mutation Rate
Time
Mutation Rate
Time

**Supplement 5b to Fig. 5C-D**

*Experiment2*:

| | |
|---|---|
| Population size | 10000 |
| Number of mutation rate genes | 1 |
| Number of adaptive trait genes | 10 |
| Inherited variance | 0.05 |
| Recombination rate | 0.05 |
| Environmental change rate | 0.001 |
| Mode of selection (stabilizing vs directional) | directional |
| Cost of mutation rate | 0.001 |

| Directional selection on adaptive trait | Directional selection on adaptive trait at monogenic inheritance of mutation rate |
|---|---|
| 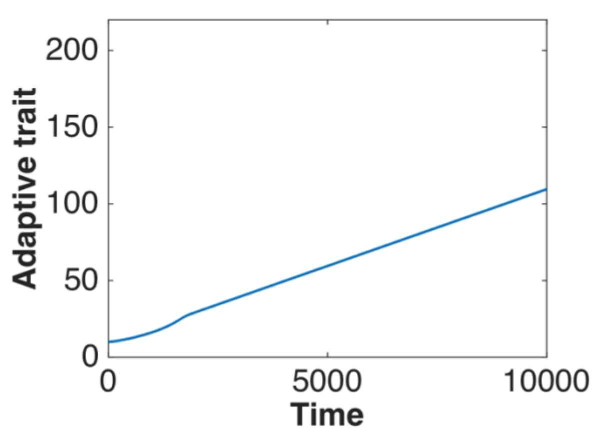<br> | 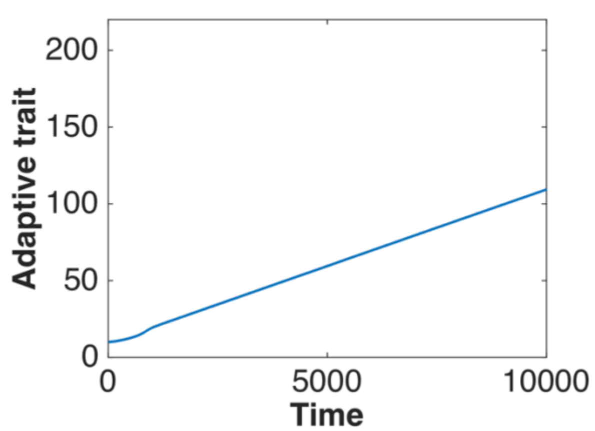<br> |
| 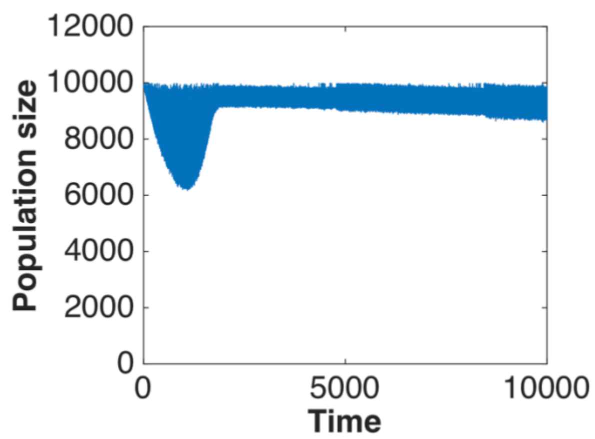<br> | 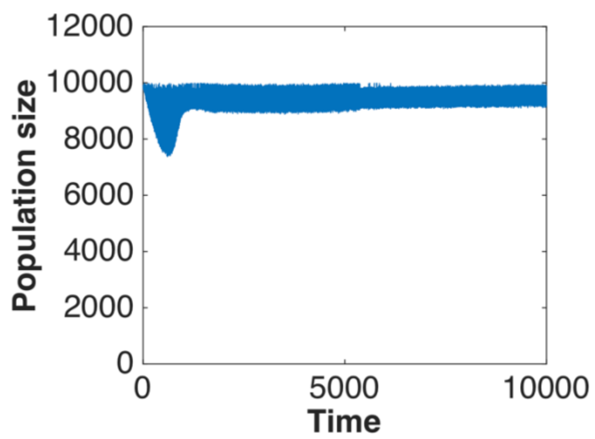<br> |

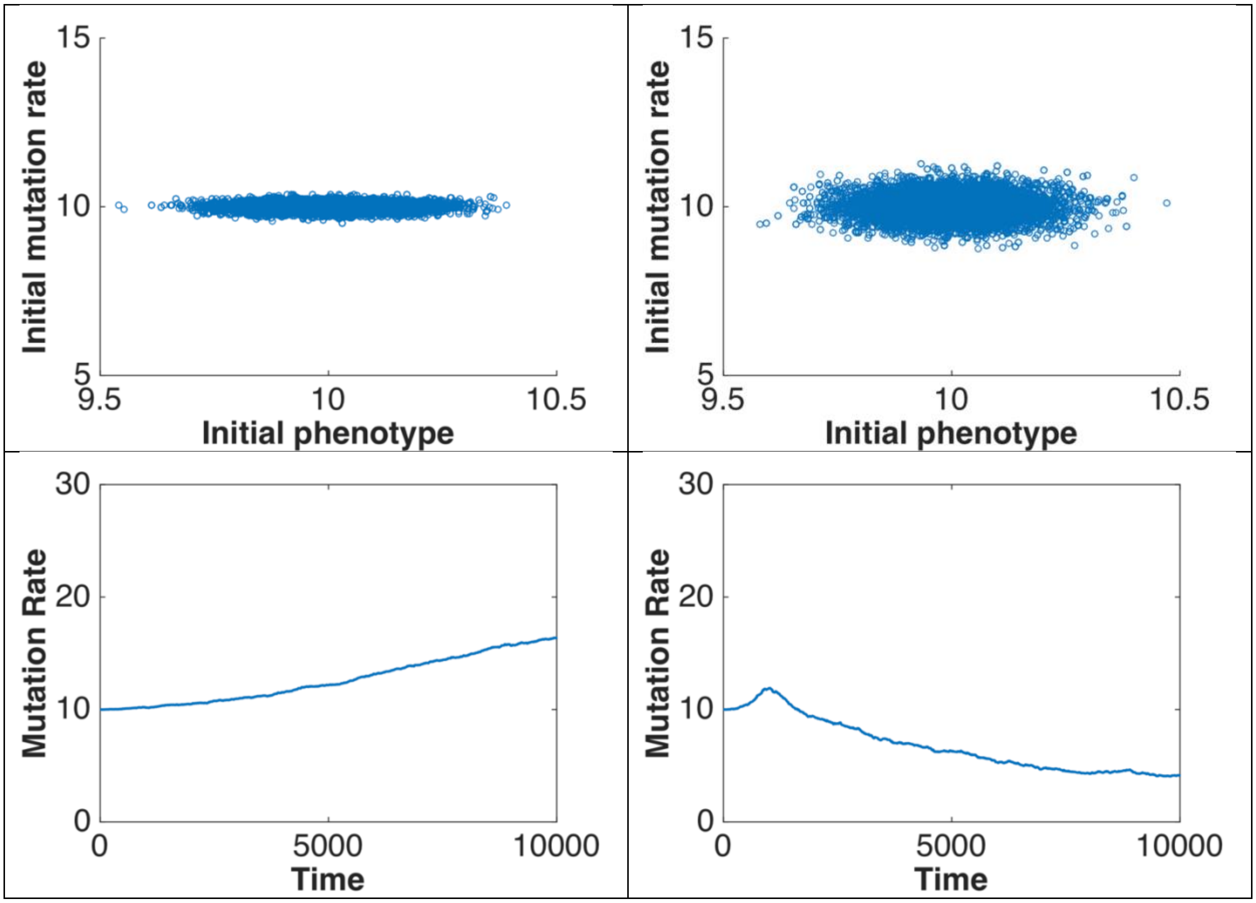

Initial mutation rate
Initial phenotype
Initial mutation rate
Initial phenotype
Mutation Rate
Time
Mutation Rate
Time

**Supplement 6a to Fig. 6A-B**

*Parameters*:

| | |
|---|---|
| Population size | 10000 |
| Number of mutation rate genes | 10 |
| Number of adaptive trait genes | 10 |
| Inherited variance | 0.05 |
| Recombination rate | 0 |
| Environmental change rate | 0.001 |
| Mode of selection (stabilizing vs directional) | stabilizing |
| Cost of mutation rate | 0.001 |

<table>
<tr><td>Stabilizing selection on adaptive trait</td><td>Stabilizing selection on adaptive trait with absent recombination</td></tr>
<tr><td>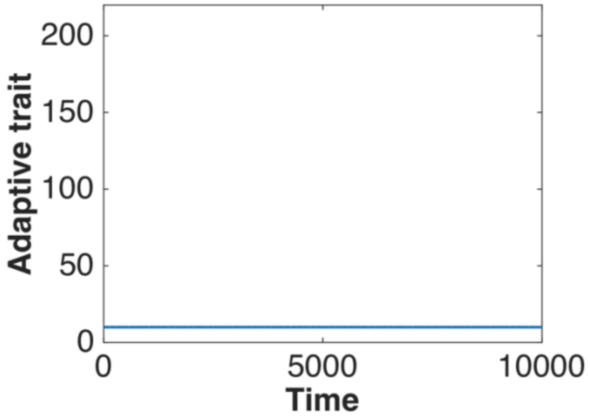
</td><td>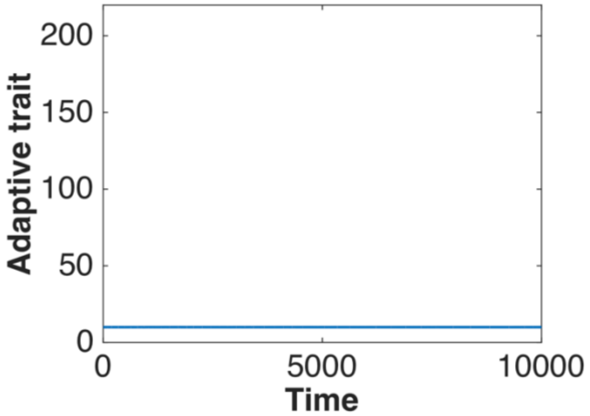
</td></tr>
<tr><td>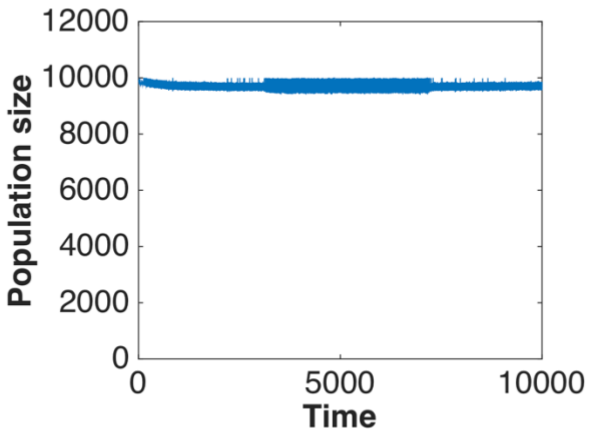
</td><td>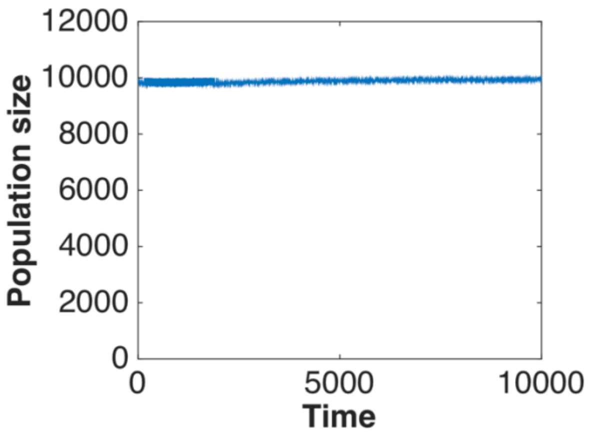
</td></tr>
</table>

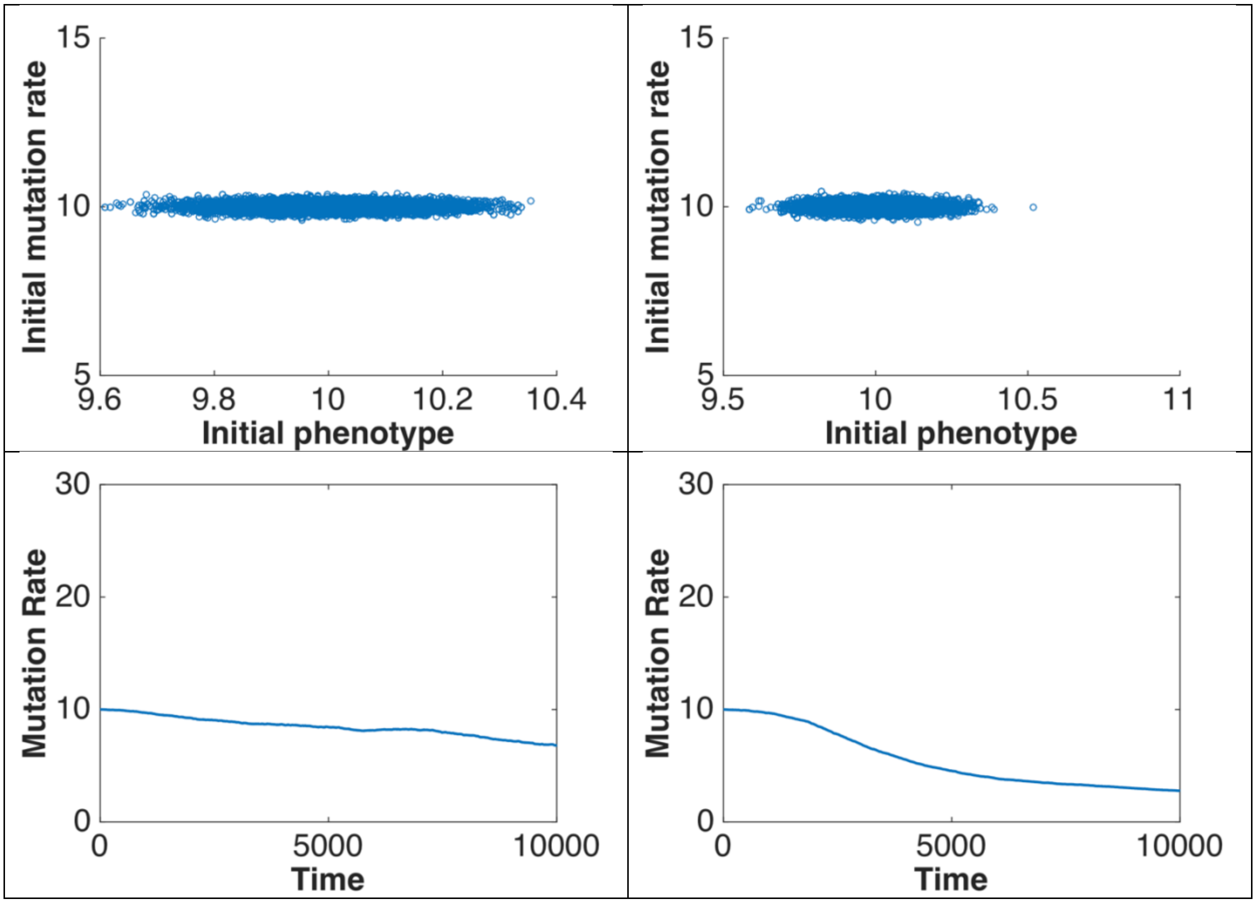

Initial mutation rate
Initial phenotype
Mutation Rate
Time
15
10
5
9.6
9.8
10
10.2
10.4
9.5
10.5
11
30
20
0
5000
10000

**Supplement 6b to Fig. 6C-D**

*Parameters:*

| | |
|---|---|
| Population size | 10000 |
| Number of mutation rate genes | 10 |
| Number of adaptive trait genes | 10 |
| Inherited variance | 0.05 |
| Recombination rate | 0 |
| Environmental change rate | 0.001 |
| Mode of selection (stabilizing vs directional) | directional |
| Cost of mutation rate | 0.001 |

<table>
<tr><td>Directional selection on adaptive trait</td><td>Directional selection on adaptive trait with absent recombination</td></tr>
<tr><td>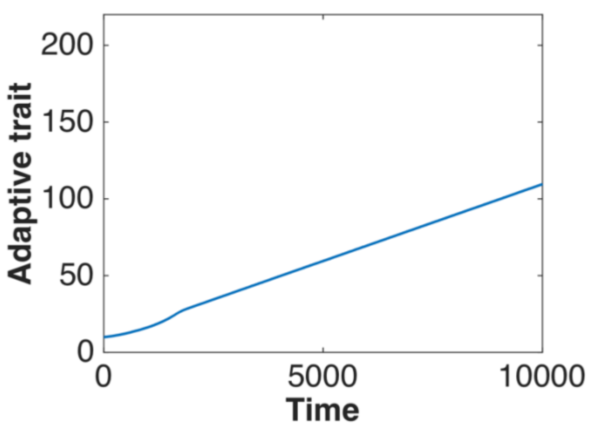
</td><td>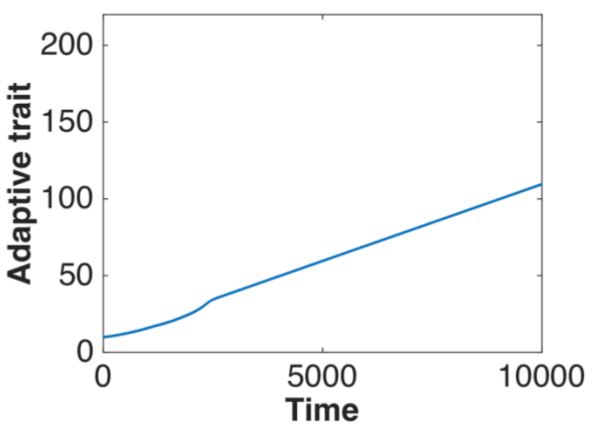
</td></tr>
<tr><td>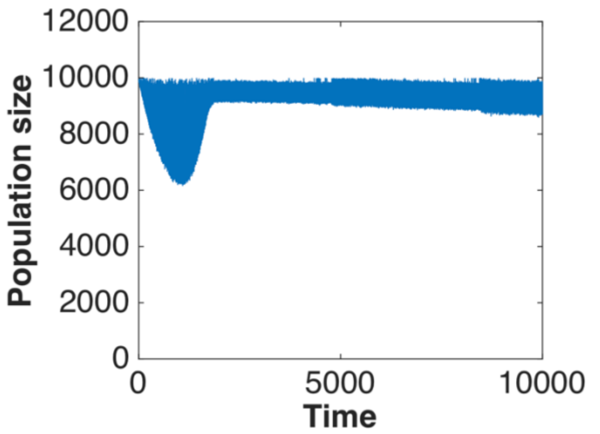
</td><td>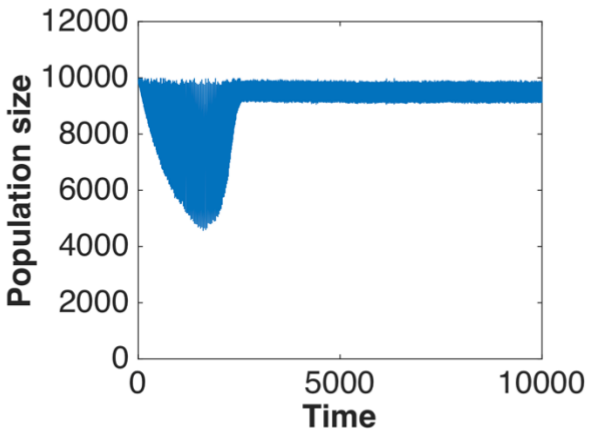
</td></tr>
</table>

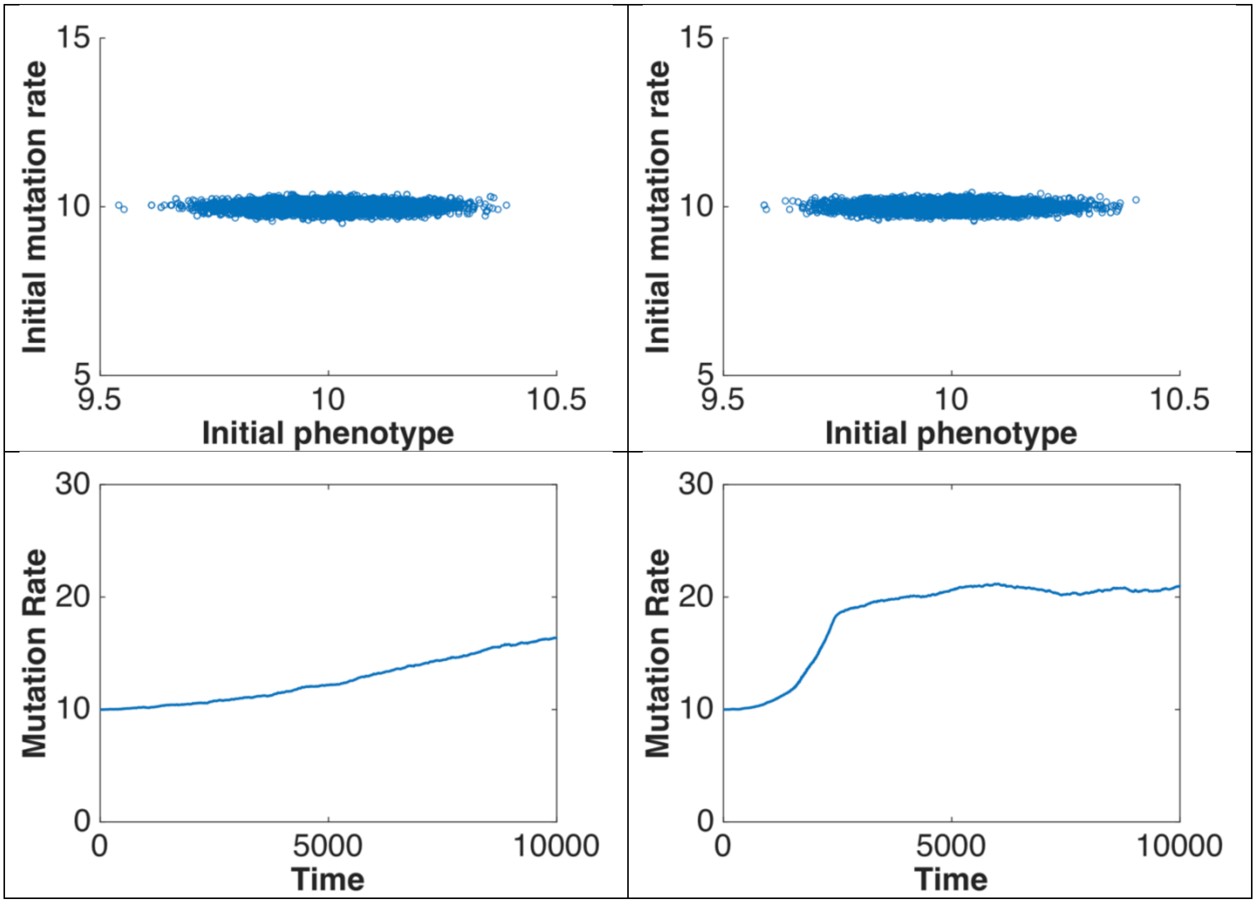
Initial mutation rate
15
10
5
9.5
10
10.5
Initial phenotype
Initial mutation rate
15
10
5
9.5
10
10.5
Initial phenotype
Mutation Rate
30
20
10
0
0
5000
10000
Time
Mutation Rate
30
20
10
0
0
5000
10000
Time
